# An information theoretic approach to community detection in dense cortical networks reveals a nested hierarchy

**DOI:** 10.1101/2024.08.07.606907

**Authors:** Jorge Martinez Armas, Kenneth Knoblauch, Henry Kennedy, Zoltan Toroczkai

## Abstract

Interareal structural connectivity explored by tract-tracing reveals super-dense, weighted, directed, and spatially embedded complex networks. These properties make the extraction and alignment of the community structure of these networks with brain function challenging. Here, we overcome these difficulties using a principled, information theoretic approach that distinguishes connectivity profiles using the Hellinger distance. Applying it to retrograde tract-tracing data in the macaque, we show that the cortex at the interareal level is organized into a nested hierarchy of link-communities alongside a node-community hierarchy. We find that the ½-Rényi divergence of connection profiles, a non-linear transform of the Hellinger metric, follows a Weibull-like distribution and scales linearly with interareal distances, establishing a quantitative expression between functional organization and cortical geometry. Finally, we show that the loop entropy along the hierarchy is maximized at a community description level that is neither excessively detailed, nor non-specific, defining a “Goldilocks” level that we hypothesize, is optimal for system-wide efficient information processing.

**Author Summary:** Understanding how functional organization emerges from connectivity alone is a central challenge in neuroscience. The cortex forms a dense network, making it difficult to identify meaningful groups of areas using standard methods. Here, we introduce a principled information-theoretical approach that compares the connectivity patterns of brain areas to uncover their community structure and organization. By first detecting the link communities and then deriving the grouping of areas from it, our method reveals a hierarchical structure based on patterns of reciprocal communication. Applied to the retrograde tract-tracing database of the macaque cortical network, this approach identifies communities that share similar interaction patterns, offering new insights into large-scale brain organization. More broadly, it provides a general framework for analyzing complex networks with dense, directed, and weighted connections.

---

**U**nderstanding how the brain works is an inverse problem where observational data is exploited to back-engineer functionality and principles of information processing. Given the brain is a complex networked system, one approach is to study how the structure of this network deviates from that of other, random networks, serving as null models. The deviations and their heterogeneity across scales reflect structural and functional specificities of the brain (Bassett & Sporns, 2017; Bullmore & Sporns, 2009; He & Evans, 2010; Sporns et al., 2005; Váša & Mišić, 2022).

A heavily used method in studying a network’s structural organization is identifying its communities (Fortunato, 2010). In their original definition, communities (or clusters) are subsets of nodes that are more interconnected amongst themselves than with the rest of the network. A more general definition considers communities to be subsets of network objects (nodes, edges, subgraphs, etc.) that are “closer” to one another by a pre-defined *similarity* measure, than to the rest of the network. When the “network objects” are themselves communities, we are dealing with communities of communities, that is, a hierarchical organization of the network (Clauset et al., 2008; Markov, Vezoli, et al., 2014; Ravasz et al., 2002; Ravasz, 2009).

Communities in network systems have been associated with components that perform specific functions (Fortunato, 2010; Henry et al., 2016; K. Li & Pang, 2013; Mesulam, 1990; Onnela et al., 2012; Ruan et al., 2010; Strong et al., 1984). Compared to homogeneous networks, heterogeneous networks have an increased encoding capacity (Markov, Ercsey-Ravasz, Van Essen, et al., 2013; Pessoa, 2014; Sporns, 2013; Sporns & Betzel, 2016; Wu et al., 2011). Since in the brain information is encoded by the activities of distributed neuronal populations (Buzsáki, 2010; Felleman & Van Essen, 1991), the network structure has a critical impact on the specificity and tunability of those activities (Michalareas et al., 2016; Vezoli et al., 2021). The extent to which a function performing network differs from a homogeneous random network is via the existence of communities, we posit that these communities constitute a key organizational feature of the brain. Further, we hypothesize that this property holds across multiple scales in the brain, including the mesoscale, where nodes represent functional areas and links represent physical connectivity in form of axonal bundles, forming the so-called interareal network.

Here, we present a novel method for analyzing the community structure of neuronal networks to help better understand the role of the connections and their weights (axon bundle densities) in shaping the network’s functions. We focus on the interareal network, given the datasets available to us, but our methodology is applicable at any scale. Due to the fact that mammalian interareal networks as exemplified by rodents and primates area are dense, directed and weighted graphs (Gămănuţ et al., 2018; Harris et al., 2019; Horvát et al., 2016; Markov, Ercsey-Ravasz, et al., 2014a), most community detection algorithms provide weakly contrasting signals. This is due to their similarity measure depending on local network density, which likely have a narrow range of variability in dense networks (Newman, 2004; Palla *et al*., 2005; Rosvall, Axelsson and Bergstrom, 2009; Ahn, Bagrow and Lehmann, 2010; Bodlaj and Batagelj, 2015). Moreover, we want to develop community definitions related to the *roles* performed by these structures so that the detected communities and hierarchies directly reflect information processing. It is important to mention that the detection of communities and their hierarchical organization is not unique, as it strongly depends on the definition of the similarity measure and on the community merging criteria. For example, in a social network, communities obtained by physical co-location (families) are not the same as those obtained by professional memberships.

Our approach is to first find the communities formed by the links/edges (Ahn et al., 2010b), and from that, to generate the node-communities and capture their hierarchical organization (if any). This follows from the assumption that links connecting similarly structured network neighborhoods share similar roles, given that the in the inter-areal network, structure is a major aspect of the algorithm itself (Horvát et al., 2016; Laydevant et al., 2024; Markov, Ercsey-Ravasz, Van Essen, et al., 2013; Molnár et al., 2024). Link based communities are also advantageous in dense graphs, given there are many more links than nodes (hence potentially more structural information). In the macaque dataset (**Figure 1a)** the edge-complete part of the network (i.e., where we have full knowledge concerning links - see Technical terms), has only 40 nodes but 999 directed edges, shown as a graph in **Figure 1b** with an approximate 3D spatial layout.

**Figure 1.**
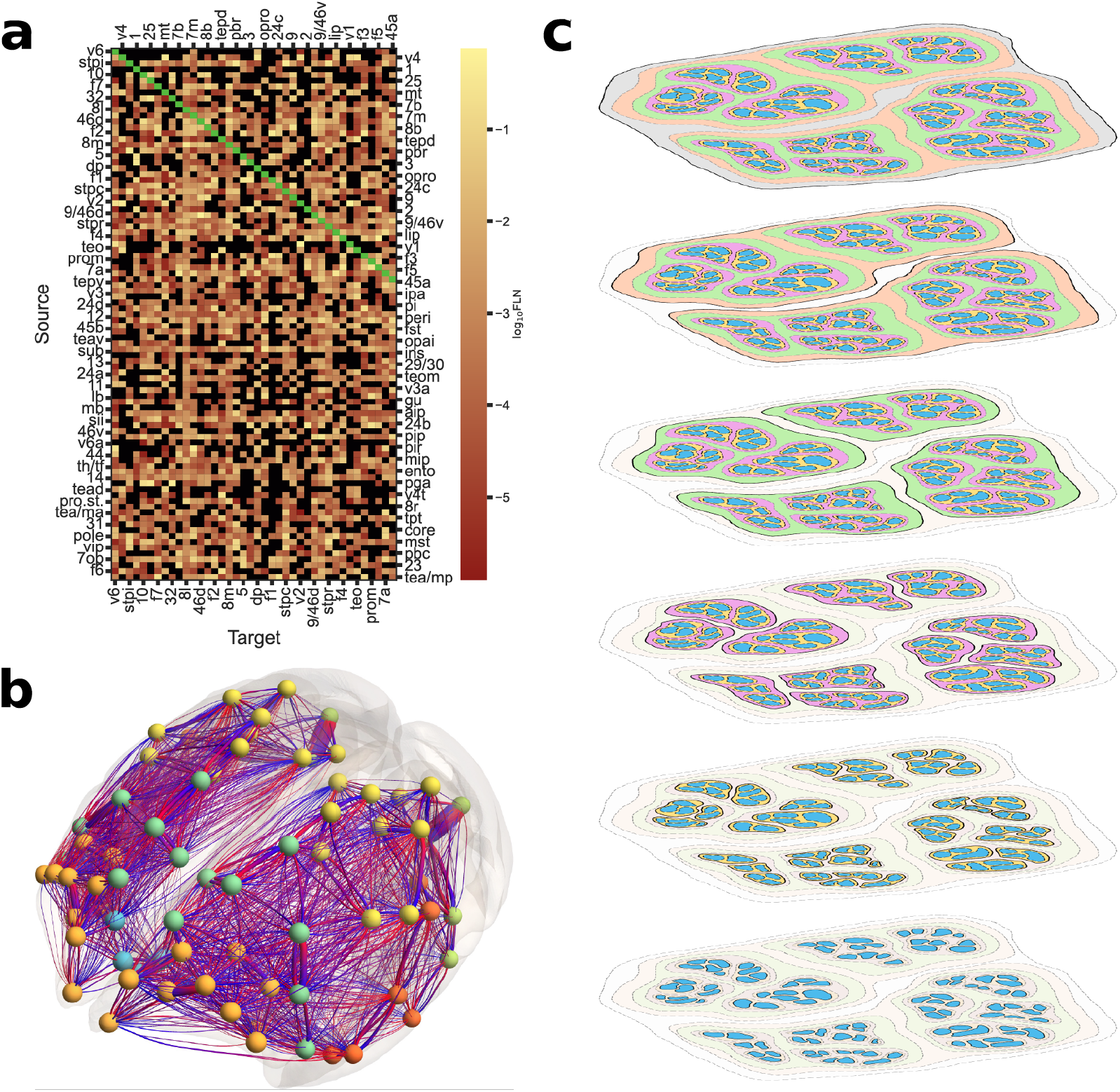
The network community hierarchy. **a**, The matrix of connection weights 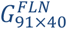 (in random order) obtained retrogradely from 40 target area injections, on an atlas of 91 areas in the macaque monkey. Weights are represented by colors, as shown in the color bar, with black and green denoting absent and intrinsic connections, respectively. **b**, The edge-complete network part 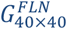 of the data from **a** represented as a network with approximate 3D layout of the area’s center of masses (network nodes) over both hemispheres. Node colors represent different brain regions. Cylindrical tubes are connections, whose blue-red color gradient and widths depict the direction from source area to target area and weight, respectively. **c**, A schematic of the nested network community hierarchy structure (edges are not shown) studied here.

After generating link communities and determining their hierarchical organization, we define the corresponding node communities and show that they, too are organized hierarchically. Given that nodes interact through links, the community structure at the level of links will induce a partitioning of the network nodes into groups. However, the extraction of node communities from link communities has not yet been proposed to the best of our knowledge. Note, here the concept of hierarchy differs from the standard view of areal hierarchy (Felleman and Van Essen, 1991), later expanded and called the “SLN-hierarchy” by Barone et al. (Barone et al., 2000), using the feedforward (FF) and feedback (FB) classification of connections. While the SLN-based hierarchy is sequential, here hierarchy is defined as that of “nested” communities, through the relationship of set inclusion, as illustrated in **Figure 1c**. In mathematics, this is referred to as compositional containment hierarchy, which establishes an order among the communities along the hierarchy depth, but there is no similar ordering among the elements contained within any given level. Some link community papers have used a binary (unweighted, undirected) connectome obtained from dMRI tractography (de Reus et al., 2014), but do not identify node communities or hierarchies. An alternative approach used fMRI functional connectivity data (Meunier et al., 2010), to locate a hierarchical organization in human brain functional networks.

In the following we first discuss the principles and criteria behind our approach and introduce a novel dissimilarity measure based on the so-called Hellinger distance (Cam & Yang, 2000; Hellinger, 1909), a measure of distinguishability of two probability distributions. This allows us to use an information theoretic measure, the ½-Rényi divergence *D*_½_, related to the Hellinger distance via a non-linear transformation, to provide an explicit relationship between the geometrical organization of the neuronal network and its functional properties. Note, our approach is generally applicable to all networks in which link weights have a probabilistic interpretation. Next, we generate the link-community hierarchy using the Ahn-Bagrow-Lehmann (ABL) algorithm (Ahn, Bagrow and Lehmann, 2010) based on the Hellinger distance as similarity measure and compare it to a null-model showing that there is significant levels of structural heterogeneity and specificity in the original brain data. We then extract a nodal level community hierarchy for the macaque cortex, as one of our main results. Finally, we propose a method that selects an “optimal”, Goldilocks node-community hierarchy using the notion of loop entropy .

## Results

### A network interpretation of function

Before developing our community detection method, we need to briefly consider the notion of function. Function within a biological (or sociological) system usually implies a well-defined and repeatable outcome in the system’s dynamics and is thus connected to the notion of order. While we can quantify disorder with various notions of entropy, we do not have a systematic quantification of order, as it can be of many different types. Here we propose a practical approach to the notion of function, by noting that function, fundamentally, is a *relationship* attribute, that is of the interactions between two or more objects. Consider, for example, the myoglobin molecule. It binds, stores and releases oxygen molecules, which on its own is a useless property. However, it becomes highly beneficial in oxygen storage and transport in the body, thus attaining its function via interactions within the muscle tissue. Note, that when specifying this function, we use the sequence: myoglobin **→** regulation of *O*_2_ and *NO* **→** muscle tissue contraction (Kamga et al., 2012). Thus, using network language, we propose that function is a property of an ordered pair of nodes (*i, j*), or equivalently, that node *i* performs an action towards node *j*. Regarding nodes as input/output units, the activity of node *i* towards node *j* serves as the input for the activity of *j*. In the brain, this is information or influence transfer (modulation). Here, we quantify the functional role of a node *i* in the network by the list of its outgoing connections and their properties, i.e., by its out-link profile. The list of incoming links to node *i*, or its in-link profile, represents the functional feedback of the *whole network* onto node *i*. In our community detection approach, we exploit both types of link profiles.

In the brain, we expect that most connections have a sustained role/function, and thus, the network of interareal connections is a direct reflection of the functional organization of the brain at the areal (mesoscopic) level. The links transfer information or modulation, which changes the state of the receiving (downstream) nodes, which in turn modulates/changes their output downstream towards other nodes, and so on.

The combined activities of groups of nodes represent diverse functions, possibly more complex/simple, depending on the higher/lower degree of connectivity of that group with the rest of the network. This property allows us to track the function-based hierarchy of the cortical network. While this network interpretation of function does not provide a translation between connectivity and behavior (which would require characterization of the nodal states and their dynamics), it is self-consistent and can be analyzed/quantified and compared across species provided the relevant mapping is available between the sets of nodes of the neuronal networks of the two species.

### Hellinger distance as connection distinguishability measure

The clusters resulting from community detection reflect the network’s organization with respect to a particular clustering criterium, formulated via a similarity measure. This reveals if two or more network elements (nodes, links, subgraphs) belong to the same cluster. As argued above, links represent function; hence, we need to define a (dis)similarity or distance measure between them. Two directed links are *neighbor links* if they share either a common starting node but different end-nodes (called out-neighbor links) or share a common end-node but different starting nodes (in-neighbor links). **Figure 2a** shows two neighbor out-links (*i, j*) and (*i, k*), and two neighbor in-links (*j, i*) and (*k, i*). All other pairs of links are considered disjoint, see **Figure 2b**. Only neighbor links can have non-zero similarity; disjointed links have zero similarity. We define the amount of similarity between two neighbor links by the *node-similarity* of their un-common nodes (*j* and *k*). In turn, we define the similarity of two nodes (*j* and *k*) via an overlap measure (defined below) between their connectivity profiles, i.e., with respect to all their weighted connections (including connections from/to the common node *i*). Specifically, as shown in **Figure 2c** left image, the similarity of the two out-neighbor links of *i*, namely (*i, j*) and (*i, k*) is given by the overlap measure between the *in-coming* neighborhoods of nodes *j* and *k* respectively, because the two out-links (*i, j*) and (*i, k*) of *i* are part of the in-link neighborhoods of *j* and *k*. If the opposite convention were chosen (e.g., comparing the out-neighborhoods of *j* and *k* for the out-links (*i, j*) and (*i, k*)), then the reference links themselves would not be included in the comparison, artificially reducing their measured similarity. Likewise, the similarity between the in-neighbor links (*j, i*) and (*k, i*) is given by the overlap measure between the out-neighborhoods of *j* and *k*, see the right image in **Figure 2c**.

**Figure 2.**
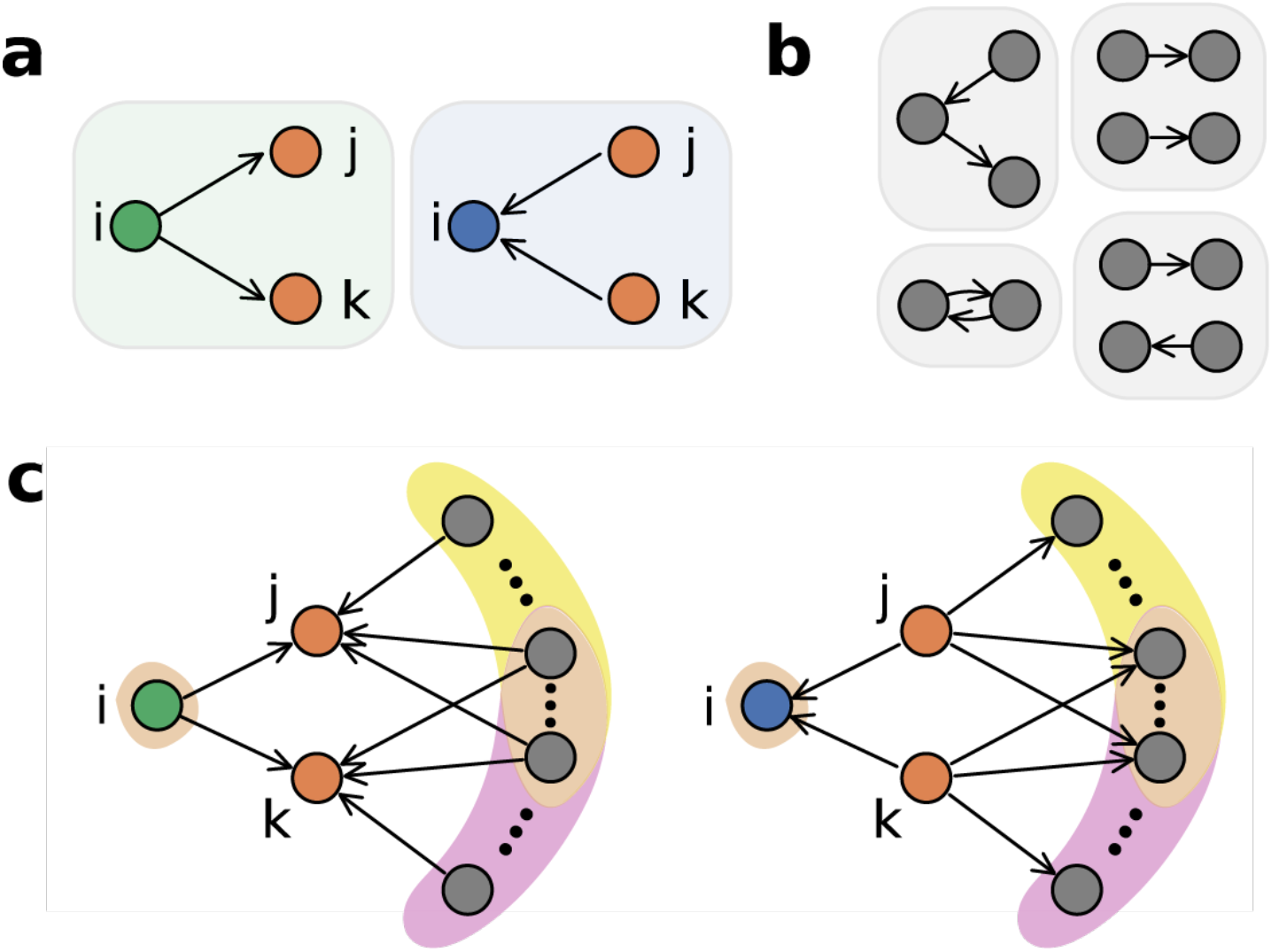
Defining the link similarity measure. **a**, We define “neighbor links” as two edges that are either both out-links or in-links from/to a common node. **b**, All other pairs of links are called “disjoint”. **c**, Similarity between two neighbor links is determined by the overlap between their weighted connectivity profiles of their non-common nodes *j* and *k*, using the Hellinger distance (see Methods section).

Nodes in a network are distinguished by their features such as their in- and out-going links and the weights on those links. By measuring feature (dis)similarities, we can embed nodes into a manifold that connects relative positions with structural likeness. Unfortunately, there are infinite ways to define such measures; however, guided by Occam’s razor principle we can formulate a rule that provides insights that other methods miss. Specifically, here we use the Hellinger distance (Hellinger, 1909), since, as shown below, it is a measure of information processing, naturally suited for our goals.

The Hellinger distance *H*(*P, Q*), used for example in nonparametric density estimation, is a measure of the dissimilarity of two probability distributions *P* and *Q*, see Methods. For our purposes it has several advantages: *i)* it is a true metric, i.e., it is non-negative, symmetric, obeys the triangle inequality, i.e., *H*(*P, Q*) + *H*(*Q, R*) ≥ *H*(*P, R*), for any *P, Q, R* probability distributions, *ii)* it is a bounded metric with 0 ≤ *H*(*P, Q*) ≤ 1, for any *P, Q* and *iii)* it relates to a practical statistical measure, namely the *sample complexity* 𝒩^δ^ (*P, Q*). A key consequence of *i)* and *ii)* is mathematical consistency for comparisons. For example, if distributions *P* and *Q* are far but *Q* and *R* are close, then it follows that *P* and *R* are also far - a property not guaranteed in general for similarity measures used in the literature. In our case, we use the Fraction of Labeled Neurons (FLN) as probability distributions, which are the link weights in the interareal network datasets. These are obtained from retrograde tract-tracing injections into the cortex, see the Methods section. The FLN weights admit a probabilistic interpretation (Markov, Ercsey-Ravasz, Van Essen, et al., 2013) so that the Hellinger distance provides a good dissimilarity measure in our context. As similarity, we employ *S*(*P, Q*) = 1 − *H*^2^(*P, Q*), called *affinity* or the Battacharyya coefficient (Bhattacharyya, 1946) (0 ≤ *S*(*P, Q*) ≤ 1) in statistics. By definition, disjoint links have the largest possible distance (*H* = 1) and thus the lowest similarity value (*S* = 0).

The Hellinger distance is directly related to the notion of sample complexity (Canonne et al., 2019), which is a measure of how many samples we need to draw from a probability distribution (or a from a process described by that distribution), in order to start distinguishing it from an alternative distribution (see Methods). The smaller the Hellinger distance, the greater the number of samples required.

**Figure 3a** shows 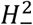 and 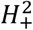 for all node pairs as a function of white-matter interareal distances between the center of mass of areas (see Methods) for the nodes within the edge-complete network (ECN). One can see another advantage of using the Hellinger distance as a dissimilarity measure: it is sensitive at all scales, especially at short distances, providing a good resolution in discernibility, hence also locally.

**Figure 3.**
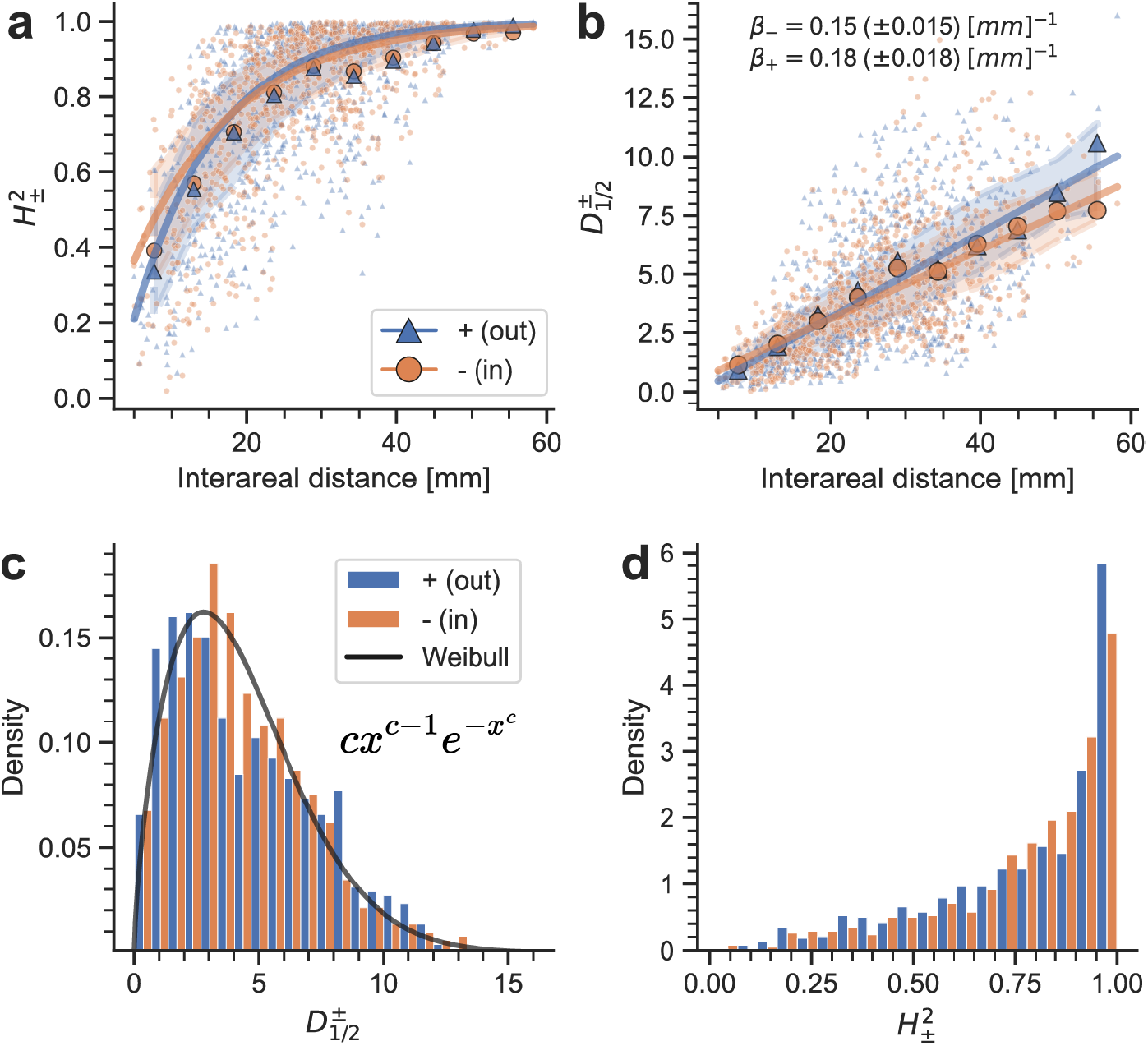
Measures of connectivity profile differences. **a**, The Hellinger distance 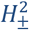 as function of interareal distance for the macaque dataset. Lines, 95% confidence level (CL) uncertainty bands, and bin averages (large markers) are derived from the linear fits in panel **b**, using equation (1). **b**, ½-Rényi divergence, an information geometric measure as function of distance, showing a linear relationship. Lines and 95% CL uncertainty bands are obtained from a weighted linear regression from ten binarized averages and inverse variance weights. Slope regression coefficients *β*_∓_ (± 95% CL) are shown at the top. **c**, The distributions of 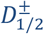 (bars) are well approximated with a Weibull distribution of shape parameter *c* = 1.67. **d**, histograms of 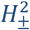.

### Rényi divergence of node-connectivity profiles

A well-established and often used statistical distance between two probability distributions (a baseline distribution and a model distribution) in information theory is the Kullback-Leibler (KL) divergence. However, the KL divergence is not suited for our purposes, given it requires a baseline distribution, and it is not symmetric, making comparisons difficult to interpret. It presents infinite divergence when the model distribution is zero and it is not a true distance metric. The Hellinger distance allows us to avoid these issues. Moreover, there is an information theoretic divergence measure that is directly related to the Hellinger distance, namely, the *α*-Rényi divergence (Renyi, 1961) which constitutes a generalization of the KL divergence. Namely, for *α* = 1/2, the ½-Rényi divergence *D*_½_ (see Methods for mathematical definition in terms of distributions *P* and *Q*) obeys the nonlinear relationship:

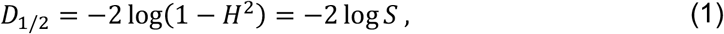

(note, *α* = 1 is the classic KL divergence) (Nielsen & Boltz, 2011). For our purposes, the ½-Rényi divergence has two main advantages: firstly, it is symmetric and secondly, it can handle zero values for any of the two probability distributions, without diverging. It has many applications in information theory, hypothesis testing, multiple source adaptation, coding and machine learning through approximate inference (Csiszar, 1995; Y. Li & Turner, 2016; Mansour et al., 2009; Morales et al., 2000; Shayevitz, 2011; van Erven & Harremos, 2014), see Methods.

Plotting the 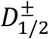 values for all ordered area pairs as a function of their distance, we obtain **Figure 3b**. Interestingly, this shows an almost linear relationship between the Rényi divergence and distances, with slightly different slopes. However, the scatter plot is heteroskedastic with significant variability for 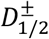 around the mean, signaling additional information. Histogram plots of 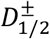 exhibit unimodal distributions with fast decaying tails (**Figure 3c**). The continuous line is a fit of the Weibull distribution 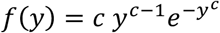 to the standardized data *y* = (*x* − *x*_*min*_)/*σ* where *x*_*min*_ is the minimum value of *x* (in our case 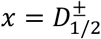 ). *c* and *σ* are referred to as shape and scale parameters and fitting gives *c* = 1.67 and *σ* = 4.75 respectively, for our macaque dataset. In the Discussion section we argue for 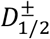 being a Weibull distribution as used in extreme value statistics (Montgomery, 2020). Note, the goodness of fit to the Weibull has an AIC index of 3554.0, whereas for the Gamma and the Lognormal distributions are 3569.4 and 3590.0, respectively. **Figure 3d** shows the histograms of 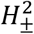 for the macaque, a monotonically increasing function, indicating that the statistics improves with increasing dissimilarity (i.e. there are more area pairs at larger dissimilarity values).

### The link-community hierarchy of the macaque cortex

Having defined a proper distance (or similarity) measure between the links, we can now use the ABL link-community detection algorithm (Ahn, Bagrow and Lehmann, 2010) on the ECN of the macaque dataset, see Supplementary Materials S3.1 for algorithm description and Supplementary Materials Figure S8.

**Figure 4a** shows the link-community hierarchy obtained with this method for the edge-complete component of our dataset. It has many intricate structures at all levels, indicating the complexity of the functional hierarchy. However, it is difficult to interpret because it essentially shows the “cabling structure between the elements of a device,” not the elements and groups of elements of the device that are working together (considered to be the natural description of the inner workings of a system). We can, however, make a comparison of the link-community hierarchy in **Figure 4a** top with the same generated from a null model, called the configurational model, see **Figure 4a** bottom, and their respective merging distance histograms. The configuration model is obtained from the original network by randomly rewiring its connections (while maintaining edge weights) (Orsini et al., 2015). The link-community hierarchy for the configuration model (conf; typical instance), in **Figure 4a** bottom, shows that the link-community hierarchy for the original macaque (MAC) is more structured.

**Figure 4.**
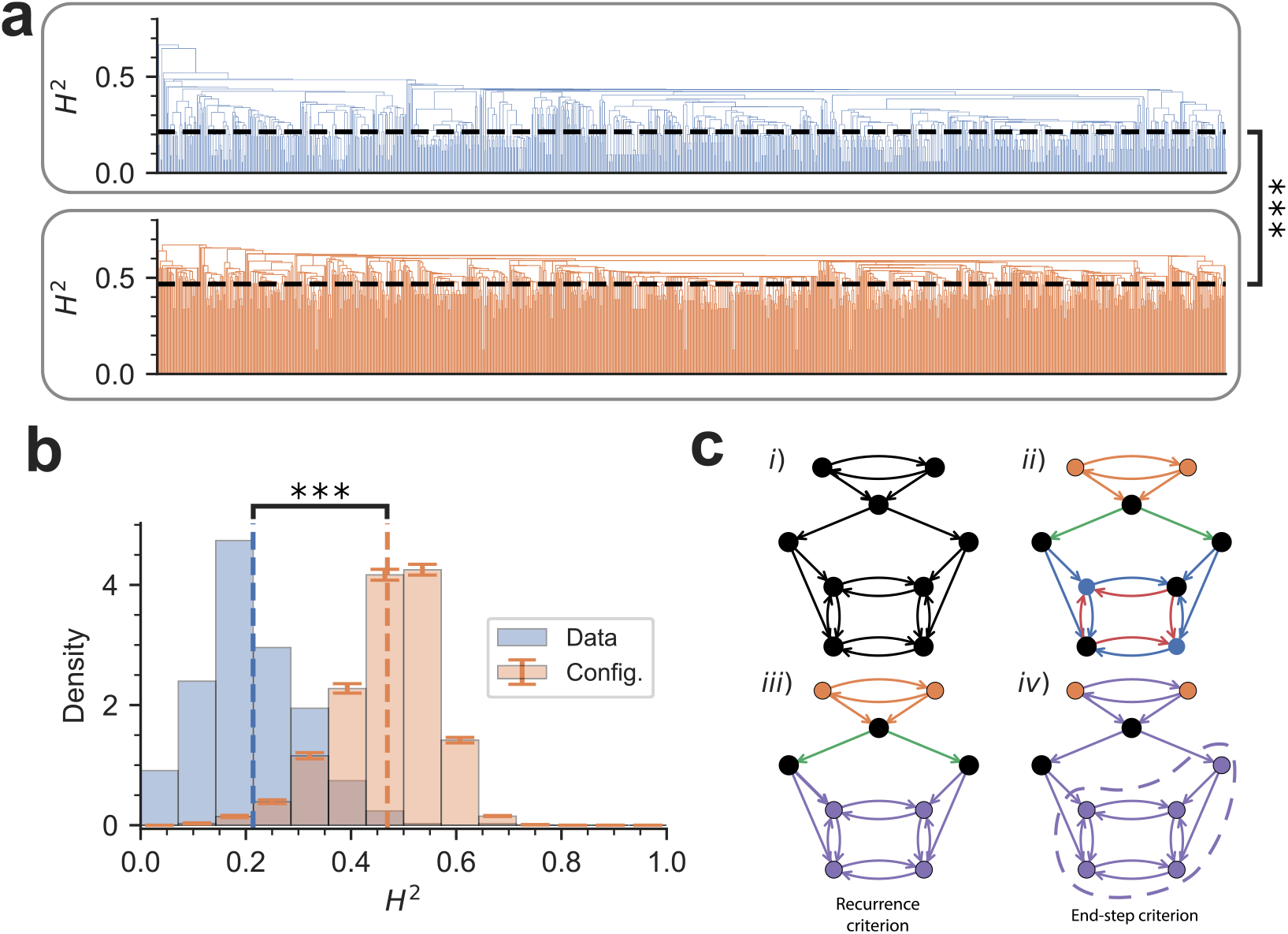
Link- and node-community hierarchies in the macaque brain. **a**, The link-community hierarchy shown as a dendrogram for the data (top, in blue) and the configuration model (bottom, in orange). The vertical lines at the lowest level represent individual links (999 links). **b**, The histograms of link community merging distances in each hierarchy. Vertical dashed lines denote mean merging (Hellinger squared) distance. The data has significantly smaller 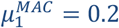 mean, compared to the configuration model’s 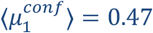 and a much larger skewness (third Fisher standardized moment), of 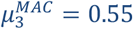, vs 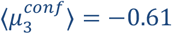 (one-sided t-tests with 1000 configuration model instances for both statistics; *P* < 10^−5^). **c**, Example of the node-community merging process. (*i*) Initial network. Black color represents single link- or node communities. Assume that in a few steps of the link merging process, described in Supplementary Materials S3.1, we obtain (*ii*), where there are 4 distinct link communities and just one node community (orange) – the black nodes have not yet joined. At the next step, the red and the blue link communities merge due to the recurrence criterion into the purple link community, and the purple node community appears. (*iv*) Finally, when all links belong to a single link community, i.e., the brown link community, the unmerged node-communities join in sequentially (end-step criterion has to be used, because the orange only feeds into the rest, no reciprocal connections).

This is also seen from the differences in the histograms of the merging events. Good quality, heterogeneously structured link communities across the whole hierarchy imply merging over small distances, indicating good separation between the distances within and between communities, from low to large distance values. In contrast, random structures will not show clear separation until large distance values, hence at a larger mean distance value. The skewness of the distribution measures where the merging rate is high or low. Positive skewness indicates the merging process has a higher merging rate at small distances (beginning of the merging process), with a rate decreasing at large distances (end of the process), consistent with a good quality hierarchical community structure. The data shows these features, whereas the randomized version (configuration model) does not, indicating that the original link community hierarchy is consistent with a pronounced hierarchical structure.

### Extracting the node-community hierarchy from the link-community hierarchy

As the link-community hierarchy is difficult to interpret, we need to generate a nodal community hierarchy derived from the link-community structure. The main criterion that we use for generating the node communities is that at a given level two communities of nodes are merged if they are bidirectionally connected with edges from *the same link community*. We refer to this as the “recurrence criterion,” i.e., the two groups must have recurrent connectivity within the same community of **Figure 4c**. However, there are node groups that cannot be connected by this criterion because, for example, they have only in-going, or out-going connections with the rest of the network; they are added into the node-community hierarchy at the end sequentially, in increasing order of their index. **Figure 5** shows the result in the form of a node community dendrogram (see Technical terms). The merging between two node communities happens at an *H*^2^ threshold value (branching point value) that is equal to the distance between two link communities associated with the two node communities (see Supplementary Materials Figure S9 for detailed explanation). The thick-colored arcs (A1–A6) denote branches of the node-community hierarchy that align with known large-scale functional systems in the cortex. It is important to note that in this representation, the separation between two areas is measured by the length of the shortest path traveling up on the tree from one of the areas, through the closest branching point that allows descending along the branches to the other area. For example, one of the largest separations is going from the primary visual area V1 (in arc A2) to area 8l of the frontal eye field (arc A6) as it requires going all the way to the branching point at the top of the tree.

**Figure 5.**
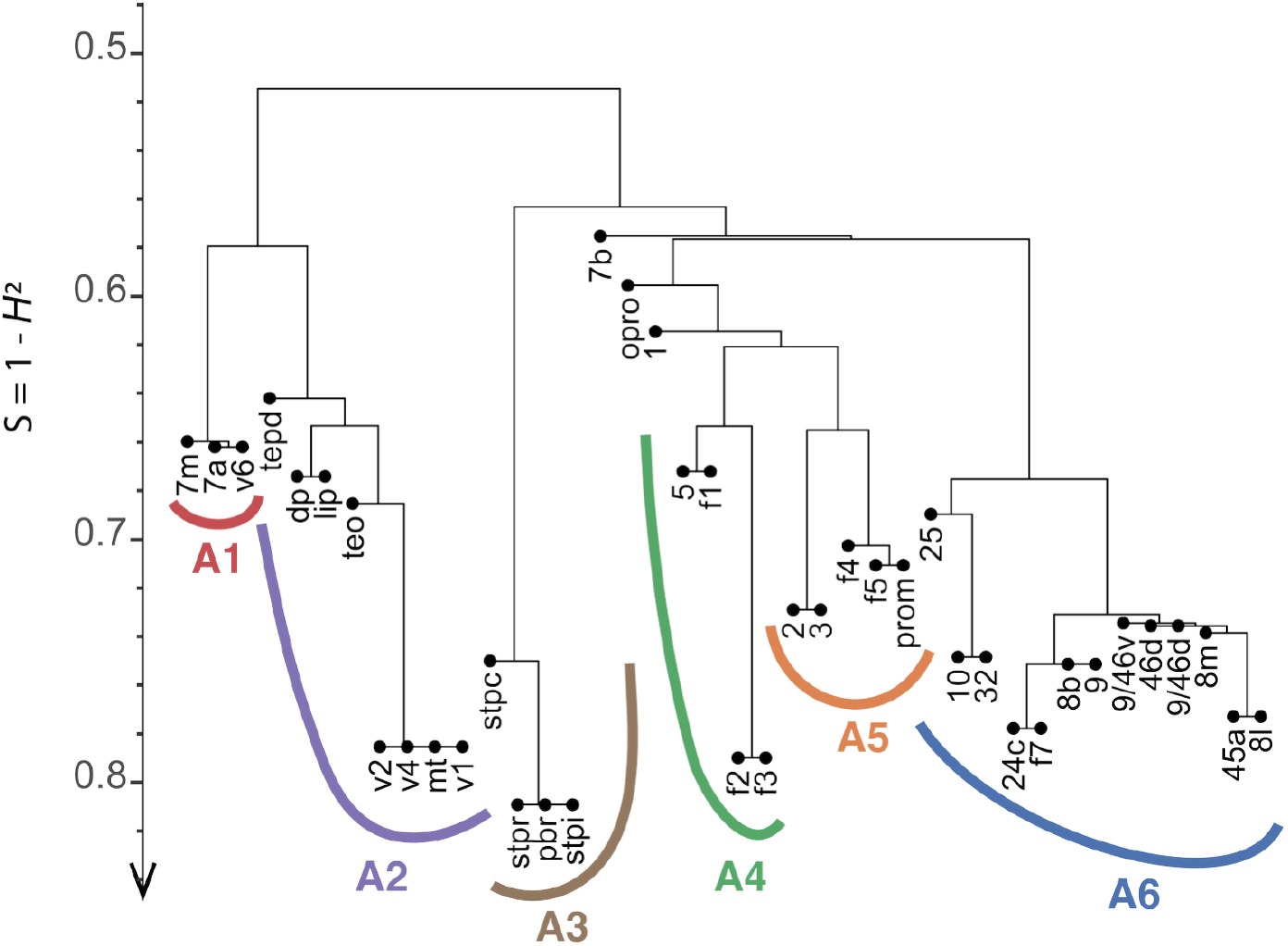
The nested node-community hierarchy in the macaque cortex at the interareal level. The y-axis is the link-community similarity between node communities at their merging point, increasing from top to bottom. Areas that merged towards the bottom (for example V1 and V2) have more similar in- and out-neighborhoods than the ones that merged close to the top (7b, OPRO, etc.). We note that areas with strong topographic organization tend to merge towards the bottom of the hierarchy (V1, V2, MT, 3, 2, etc.). The colored arcs correspond to functional brain regions preponderantly associated with movement detection (red, A1), movement/object recognition (purple, A2), multisensory (brown, A3), motor (green, A4), somatomotor (orange, A5), and high-level cognitive functions (blue, A6). The horizontal arrangement of the areas is computed with the *ggtree* software (Yu et al., 2017) and signals no special ordering.

As noted by (Hilgetag & Goulas, 2020), several types of hierarchies have been proposed in the context of mesoscale connectomes, each capturing different properties, such as connection topologies, gradients in areal features, or laminar patterns. Notably, they hypothesized the importance of a “modular” hierarchy describing the organization of the network at multiple scales. Our results are in direct support of this hypothesis and our method, to the best of our knowledge, is the first to determine the hierarchical organization in nested communities. Crucially, this hierarchy differs from others as it is not sequential, but rather a nested structure of more homogeneous groups interconnected in heterogeneous ways from the smallest to the largest scale. Although results have been published on the community structure in other mammalian brains (Harris et al., 2019), those were centered on the community partition, rather than the study of the hierarchical organization of nested groups of communities.

### A “Goldilocks” level of the node-community hierarchy

The node-community hierarchy shows the network of clusters from the most detailed level (one node in every cluster) to the least (the whole network is one cluster). The first contains too much information, but the latter is uninformative. While the hierarchy as a whole provides an overall picture of the network’s structural organization, we may ask if there is an “optimal” level that is neither too detailed, nor too uninformative, but “just right” within the hierarchy, known as a “Goldilocks” criterion (Kidd et al., 2012). Ahn, Bagrow and Lehmann, 2010 uses the average link density over all link communities to identify such a level (see Supplementary Materials S3.6), which, in the sparse network examples used in that study, has a peak in the middle of the hierarchy. However, for dense networks such as ours, this does not work, as link densities increase all the way to the top of the hierarchy (**Figure 6a**). Here, we modify the Goldilocks criterion so that it exhibits a robust peak within the hierarchy. The process we use is similar to the Ahn et al. measure, based on the number of links in excess of the tree structure within each link community, but calculates an entropy measure related to directed network loops, and chooses the hierarchy level that maximizes this “loop entropy” (see Methods). The optimal level in the link-community hierarchy is thus the one in which the distribution of the “excess” links (i.e., links generating loops and thus recurrence of information flow) is the most uniform across all link communities. The level thus defined is the most homogeneous with respect to recurrence. One can show that at either end of the link-community hierarchy *S*_*L*_ = 0, and thus it exhibits a global maximum. Given that the node-community hierarchy is derived from the link-community hierarchy, we simply use this correspondence to read off the Goldilocks level of node-community hierarchy. This shows that at the optimal level, the macaque 40×40 dataset has a total of 6 communities (and also additional three, single node-communities), see **Figure 6b-c**. This may change as the dataset grows, but the algorithm will identify the optimal level in any version of the dataset. In the Supplementary Materials Figure S18, we show that our analysis of a 29×29 dataset (Markov, Ercsey-Ravasz, et al., 2014a) results in a similar node-community hierarchical structure.

**Figure 6.**
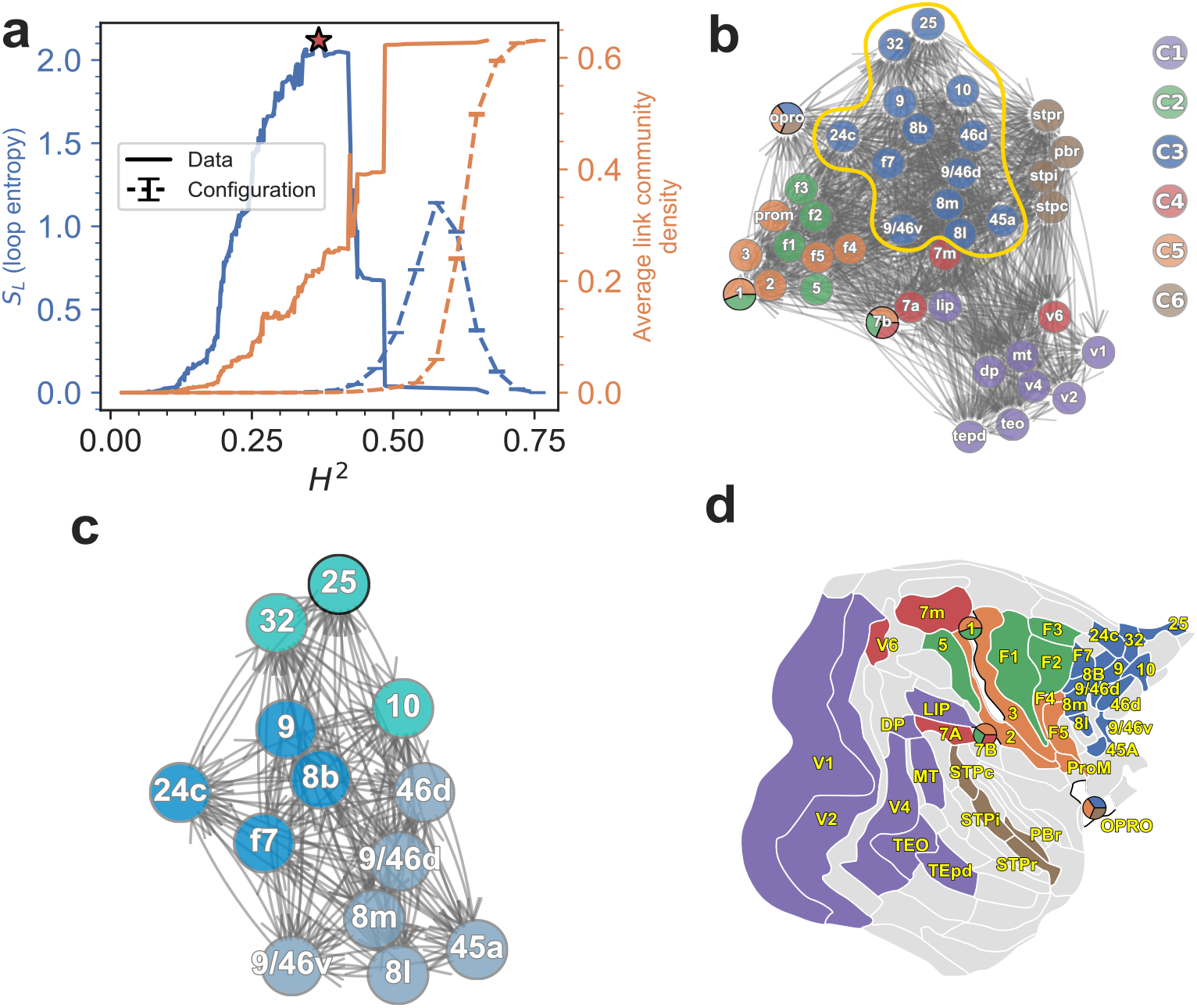
Community structure at the Goldilocks level of the node community hierarchy. **a**, Distribution of the loop entropy *S*_*L*_ (solid line) and the average LC density (Ahn, Bagrow and Lehmann, 2010) (dashed line) as function of dissimilarity *H*^2^. Solid lines correspond to the empirical network, while dashed lines show the mean trajectory of 100 configuration model instances (error bars = SEM). The red star marks the maximum loop entropy in the empirical network. The configuration model exhibits a lower maximum loop entropy than the empirical network, indicating reduced hierarchical information and weaker community diversity. **b**, The optimal node-community structure corresponding to the maximum loop entropy value (the star in **a**). Nodes with pie-charts in them represent nodes with overlapping communities (NOCs; see **Table 1** for more information about them). **c**, The sub-community structure (a few levels below) of the blue community from **b** within the yellow loop. Area 25 community membership was found subsequently, using the cover assignment algorithm (see Supplementary Materials S3.4 section). **d**, The optimal NC structure on the cortical flat map. Grey areas do not belong to the 40×40 edge-complete graph.

### Nodes with overlapping communities

The algorithm generates a small number of nodes that have multiple community memberships, i.e., nodes with overlapping communities or NOCs (**Figure 6b**). Such nodes are likely multifunctional, as shown with a pie-chart structure in **Figure 6b**. While this set will change with the growth of the data, some are likely conserved (See Supplementary Materials S3.4 for how areas are assigned to one or more communities).

**Table 1.** Cover neighborhood direction of the nodes with overlapping community membership in Figure 5b. At the optimal loop entropy level, three areas do not merge with any non-single node community. As seen in **Figure 5b**, the non-single node community ids are C1, C2, C3, C4, C5, and C6. Using the cover assignment algorithm, those areas are assigned to some non-single node communities by maximizing their average in/out-neighborhood similarity, or both (see Methods section). The neighborhood direction information is shown here.

| Area | In | Out | Both |
| --- | --- | --- | --- |
| 7b | C2 and C4 | - | C5 |
| 1 | C2 | - | C5 |
| OPRO | - | C3 and C6 | C5 |

Our method is feature-based and addresses specific challenges encountered with established community detection approaches such as modularity maximization, Infomap, and stochastic block models. These methods remain highly influential and effective in many contexts, but in the dense, weighted, and directed interareal network, they show limitations. For example, modularity maximization was unable to separate the somatosensory–premotor and premotor–motor systems (C2 vs C5), consistent with the known resolution limit (Fortunato & Barthélemy, 2007). Infomap and WSBM provided reasonable partitions at a coarse scale but did not explicitly reveal the nested hierarchical organization. By contrast, our approach yields both link- and node-level hierarchies and does so deterministically and nonparametrically. Crucially, it is grounded in the probabilistic interpretation of FLNe weights: incoming connections to each area form a probability distribution, which we compare using the Hellinger distance. This probabilistic, information-theoretic framework distinguishes our method from alternatives based on overlap or correlation measures, typically used in the social network literature (e.g., Jaccard, cosine, Pearson), which in general, produces biologically less plausible results (see Supplementary Materials S1–S2 for side-by-side comparisons).

### Model comparison

In order to quantify how much overlap there is between the community structure at the Goldilocks level of a model and that of the data, we use the notion of omega index, (Gregory, 2011), see Methods. The value of *ω* = 1 corresponds to perfect overlap between the model and data. In **Figure 7a**, we show the distribution of the omega index for two network models, namely the configuration model and the EDR model. The EDR model (Ercsey-Ravasz et al., 2013) is a generative interareal network model that uses the observed exponential decay of axonal bundle lengths with exponent *λ* (for the 40×91 dataset, *λ* = 0.16 *mm*^−1^) and the same matrix of distances between the areas. In **a**, the red vertical dashed lines represent the averages of the omega index over 1000 network samples from the respective models. The fact that ⟨*ω*⟩ = 0 for the configuration model means that the connectivity randomization completely removes the community structure found in the dataset. In the EDR model ⟨*ω*⟩ = 0.44, showing that the EDR not only captures many network properties of the data (Ercsey-Ravasz et al., 2013; Markov, Ercsey-Ravasz, Van Essen, et al., 2013) but also a portion of its community structure. In **Figure 7b**, we plot the 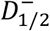 divergence for all area pairs for the two models and the data, as a function of the distance between the pairs, showing that randomization indeed completely removes the heterogeneity of the structural organization for the configuration model, whereas the EDR captures fairly well the same trend, up to medium distances.

**Figure 7.**
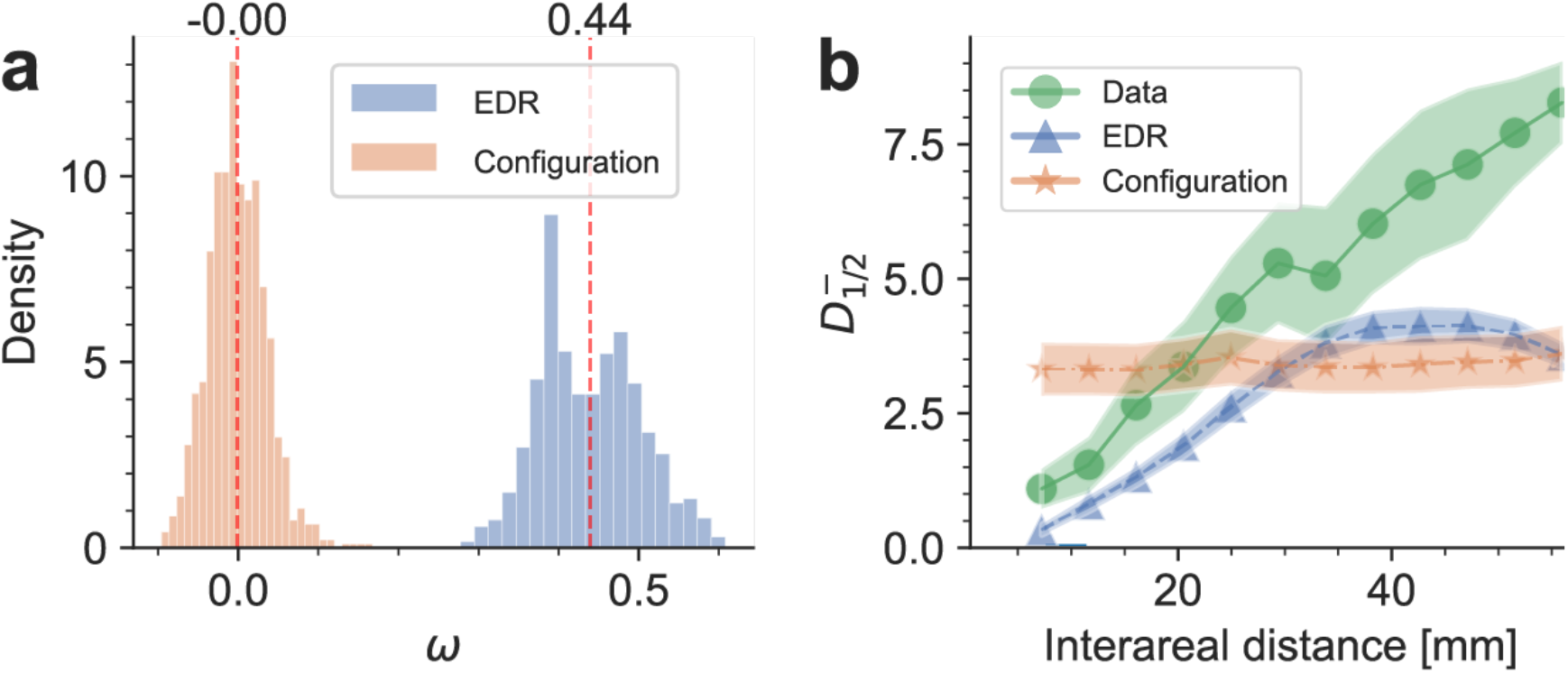
Comparison with models. **a**, Distribution of the omega index (a measure of the overall amount of overlap between two community structures; see Methods) for two network models, namely, the configuration model and the EDR model. The vertical dashed lines correspond to the average values after 1000 network samples. **b**, Similar plot as in **Figure 3b** for 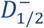 of the data and the same for the two models. Lines and markers denote the average trends. Uncertainty band widths are equal to one standard deviation.

## Discussion

Here we introduce a method to extract the community structure of the macaque interareal cortical network and its nested, hierarchical organization (compositional containment hierarchy). A chief novelty of our method is to first generate the community structure and its hierarchical organization at the level of links which allows extracting the corresponding nodal community structure and hierarchy. Given that in dense networks there are many more links than nodes, link communities provide richer information than node communities, but they are also more difficult to interpret. As the interareal network of the mammalian cortex is extremely dense, directed, and weighted, the standard community detection methods experience limitations due to using similarity measures related to local connection density, a not well-contrasting feature in ultra-dense networks. The standard methods may also neglect link directions, and/or are parameter dependent. Crucially, most methods do not reveal the hierarchical organization of the communities.

The method introduced here can be regarded as principled in a specific sense: it builds directly on the probabilistic interpretation of interareal connection weights and applies the Hellinger metric, a well-defined information-theoretic distance, to compare connectivity profiles. This embeds the network into a metric space, enabling consistent comparisons and a natural hierarchical depth scale. Because link weights (axonal bundle strengths) can be interpreted as probabilities, the Hellinger metric (a symmetric form of information theoretic divergence measure) provides a rigorous procedure for defining distances between nodes and links. This interpretation applies to any network where link weights have a probabilistic significance. In our case, weights are fractions of labeled neurons (FLNs) obtained from retrograde tract-tracing experiments, which serve as proxies for communication bandwidth between functional areas (Markov, Ercsey-Ravasz, Van Essen, et al., 2013). The relation of the Hellinger metric to the information-theoretic ½-Rényi divergence serves as a measure for the statistical distinguishability of areal neighborhoods and, interestingly, linear scales with physical distances. This further corroborates previous observations that connection profiles diverge with spatial separation, on average (Markov, Ercsey-Ravasz, Lamy, et al., 2013). Finally, we observed that the ½-Rényi divergence is distributed according to a Weibull distribution, with extreme value statistics. The same distribution was found to describe shortest path distributions in random networks (Bauckhage, Kersting, Rastergahpanah, 2013). Note, using the physical distance between the areas as an alternative to the Hellinger measure, does not work well (see Supplementary Materials S3.7).

Function in the brain is associated with links weights between areas that process similar information. When extracting the link communities (Ahn, Bagrow and Lehmann, 2010), the sequential link merging process first merges those links that perform very similar functions, and then gradually merges those link communities that are increasingly different (but can be merged at that level). At this stage the loop entropy *S*_*L*_ (**Figure 6a**) starts to increase, meaning that recurrence between the nodes of the link community also increases and the information flow structures become both more global and more complex, likely playing the role of information integration across computational modalities. Recurrence allows fast and reciprocal coordination between different functional modalities, for example, generating a fast motility response to a stimulus involving both visual and auditory streams. Thus, when creating the node communities from the link-community hierarchy we chose a merging criterion that demands recurrence between two node communities with links that belong to a given functional modality (i.e., to a given link community within that level). As for the Goldilocks level of the node-community hierarchy, we choose the one in which recurrence is the most uniformly distributed (maximum of loop entropy), being a good representative of an operational community structure. The reason for this is as follows. At the lowest levels of the hierarchy processing is localized, mechanical and non-contextual. Going up in the hierarchy, there is more communication between the different processing modalities, and the appearance of contextual embedding. We speculate that the Goldilocks level is where there is sufficient contextualization and meaning for efficient and optimum information processing across the whole network (maximum loop entropy), ideal, for example, for fast decision making, in line with the global workspace hypothesis (Dehaene et al., 1998). We also argue that working with loops and loop entropy is key, because loops form recurrent information exchange structures, required for processing information, as opposed to just unidirectional connections between two communities (e.g., from A to B) corresponding to mere information transfer. Our findings directly support the hypothesis that the brain’s hierarchically modular structure maximizes the flexibility of the brain while maintaining stable critical-like dynamics (Hilgetag & Goulas, 2020).

Supplementary Materials Figure S11 compares a randomly re-ordered ECN matrix with the one obtained from the node-community hierarchy. Supplementary Figure S12 presents the complete FLNe matrix re-ordered by node communities, highlighting that the in-neighborhoods (columns) of different communities exhibit distinct connectivity profiles. Importantly, our algorithm uses the full information of the network to compute communities, rather than being restricted to the ECN, typically considered by conventional community detection methods. For macaque, the Goldilocks level is at *H*^2^ ≅ 0.37, corresponding to 56 link communities (see Supplementary Materials Figure S13). The associated node-community network at this level has six communities (and three additional, single-node communities). By contrast, in the configuration model the Goldilocks level is shifted to higher values, 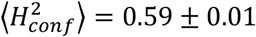 (SEM), corresponding to 77.9 ± 1.5 (SEM) link communities. Thus, the randomized model produces more numerous and mutually dissimilar link communities, in which distinction/separation between clusters is weaker than for the original data. As validation, when we applied our algorithm to sparse benchmark networks (Bryant et al., 2026) with predefined community structures, it correctly identified those communities (see Supplementary Materials Figure S14).

The resulting node-community hierarchy shown in **Figure 5**, has interesting properties. The thick colored arcs depict branches in the hierarchy as generated by the algorithm and remarkably, agree with known functional systems in the brain. Note, this is algorithm output, not “added in by hand”. For example, the branches above the A2 arc include visual areas belonging to both ventral and dorsal streams (e.g., V4 and MT), indicating that the clustering within this dataset does not strictly recover the classical “what/where” dichotomy. Rather, the grouping reflects similarities in large-scale directed connectivity patterns, as evidenced by their comparable input–output profiles in the reordered FLNe matrix (the lighter blocks around the green diagonal in Supplementary Materials Figure S12). Specifically, this hierarchy captures organization based on shared network architecture. p. The other arcs have similar interpretation, i.e., A3 includes auditory/multisensory, A4 includes motor, A5 somatomotor and A6 includes high-order prefrontal areas. Interestingly, the fact that almost all the high-level cognitive areas belong to one main branch, namely A6, suggests that areas that are not clearly associated with senses, may form their own sub-hierarchy, likely linked to attention, decision making or emotions. Our algorithm helps disentangle in a principled manner, the granular nature of functional processing in the brain. Another interesting observation is that areas with known strong topographic (e.g., retinotopic, somatotopic) organization (e.g., V1, V6, 3, 2), appear in **Figure 5** as low-level groups. Because the hierarchy is derived from similarity of directed in- and out-connectivity profiles, their placement at the bottom of their branch is consistent with these primary regions having comparatively stereotyped connectivity, whereas areas higher along the same branches show more heterogeneous connectivity profiles. A parsimonious interpretation is that topographic sensory constraints, ultimately rooted in the spatial organization of peripheral receptors and their early cortical mappings, contribute to producing structured, reproducible patterns of long-range inputs and outputs. Through successive interareal interactions, these constraints could then propagate to neighboring regions, yielding graded transitions in connectivity similarity along each branch; further testing this hypothesis will require finer parcellations (e.g., retinotopic subdivisions) and broader sampling.

Our algorithm identifies nodes with overlapping communities, which we believe play a crucial role in coordinating information flow between multiple systems. Comparisons of the node-community hierarchy with null models showed (**Figure 7**) that randomization eliminates the hierarchical organization, i.e., there is strong and specific functional organization in the cortex. The EDR model captures about 44% of the organization, meaning that the rest is due to other factors, which remain to be elucidated.

The present community detection method assumes only one node type; however, a more detailed study could employ multipartite graphs (i.e., with various node types), which could accommodate connectomes with several cell types such as in the fly (*Drosophila melanogaster*), larval zebrafish (*Danio rerio*), or different microscopic neural tissues and neurotransmitters. At the neuroscience level, we centered our attention on the macaque connectome obtained from retrograde injections; however, this dataset is incomplete, featuring 40 from 91 areas, so a community detection analysis may present misalignments with respect to the full connectome. Acknowledging this limitation, we analyzed a smaller connectome of 29 areas that was published earlier (Markov, Ercsey-Ravasz, et al., 2014b) and found that the results between the two datasets strongly agree and the general conclusions are the same.

It is important to note that there is no established ground-truth partition of cortical areas into communities, and ultimately such knowledge will require multimodal evidence encompassing function, morphology, and genetics. Our aim here is not to claim such a partition, but to provide a reproducible framework that extracts hierarchical community structure directly from the empirical connectivity in an optimized, principled manner. By grouping cortical areas according to the similarity of their connectivity profiles, our method captures an organizing principle of self-similarity, consistent with evidence that shared structural features predict both interareal connections and functional specialization (Betzel & Bassett, 2018; Beul et al., 2017; Ercsey-Ravasz et al., 2013), complementing flow-based (Infomap) and generative (SBM) approaches. In this way, the partitions we obtain should not be seen as definitive modules, but as testable hypotheses that can guide and be refined by future experiments toward uncovering biological principles of neural architecture.

## Methods

### Data

We rely on two datasets from retrograde tract-tracing experiments in the macaque monkey, obtained with consistent methods. The datasets are the matrices 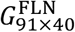 of FLN values *FLN*_*ij*_ for 40 injected target areas (*i* is a source area, projecting into target/injected area *j*), in a 91-area parcellation. The second one is the dataset that contains all pairwise distances along estimated shortest physical paths avoiding anatomical obstacles, between area barycenters, recorded in the matrix *D*_91×91_. All data is available from (Froudist-Walsh et al., 2021).

Community detection must be done in systems or subsystems with complete connectivity information. For that reason, from 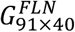, we extracted the subnetwork with 40 injected areas 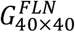 - also referred to as the edge-complete network (ECN) - whose connectivity and connection strength information have been measured between every pair of its nodes. Therefore, the communities and hierarchies are representative of the ECN, and that is not going to change when future injections are added to the dataset.

### Probability mass functions of area connections

Let us define the set of areas from the whole network as *U* and from the ECN as *V* ⊆ *U* with sizes |*U*| = 91 and |*V*| = 40, respectively. Each area *i* ∈ *V* has two neighborhoods, one for its in- and another for its out-links. The in-neighborhood of (the target) area *i* is defined as

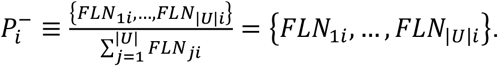

Note that this probability mass function has data from the 91 areas, with 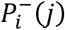 representing the probability that a neuron is projecting from source area *j* into target area *i*, with normalization 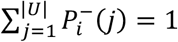, due to the definition of the FLN.

For the out-neighborhoods, the probability mass functions will cover only areas in the ECN, because an injection reveals all the inputs from the rest of the network into the target area, but not the outputs. The outputs from a given source area need to be collected across all the injections. Since there are 40/91 injections, full connectivity information (and their FLN weights) exists only for the ECN. Thus, for (the source) area *i*, it is defined as

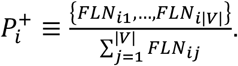

Thus, 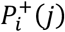 represents the probability that a neuron sampled from source area *i* projects into target area *j*, given that it projects into some area of the ECN.

### Hellinger distance

is a measure between two probability distributions (Cam & Yang, 2000; Hellinger, 1909). As explained in the main text, the Hellinger distance *H* is a metric, i.e., it satisfies the triangle inequality, and it is also bounded 0 ≤ *H* ≤ 1. For simplicity, let us first define the squared-difference function 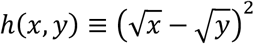. In general, the squared Hellinger distance between the probability mass functions *P* and *Q* with common discrete support *V* is defined as:

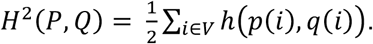

In our case the probability distributions are associated with the elements of the set of nodes *V* of the ECN, namely we are computing the squared Hellinger distance of the probability mass functions associated with nodes *A* ∈ *V* (that is *P* ≡ *P*_*A*_) and *B* ∈ *V* (*Q* ≡ *P*_*B*_) respectively, for all pairs of nodes (*A, B*). Thus, *p*_*A*_ (*i*) ∈ *P*_*A*_ represents the probability that node *i* ∈ *V* projects into (or from) node *A*. One can think of *i*’s link to (from) *A* as an (projection) “event”. The definition for *p*_*B*_(*i*) ∈ *P*_*B*_ is similar. Because the probability mass functions are associated to the elements of the support itself, the sum above needs to be computed with care when *i* = *A* or *i* = *B*, because in our dataset self-connections (intrinsic connections) are zero: *p*_*A*_ (*A*) = *p*_*B*_(*B*) = 0. To illustrate that, assume that nodes *A* and *B* have identical neighborhoods (outside of *A* and *B*), and identical probability mass functions (including *p*_*A*_ (*B*) = *p*_*B*_ (*A*)). If we were to compute *H*^2^ naïvely, element-wise, the terms *h*(*p*_*A*_ (*B*), *p*_*B*_(*B*)) and *h*(*p*_*A*_ (*A*), *p*_*B*_(*A*)) would both be non-zero, and we would obtain a non-zero squared Hellinger distance, *H*^2^ (*P*_*A*_, *P*_*B*_) ≠ 0, which would be incorrect, given that the two distributions are identical. For this reason, the correct expression of the squared Hellinger distance that we need to use is:

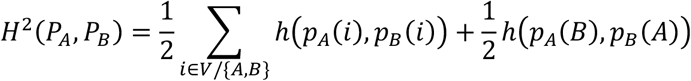

or

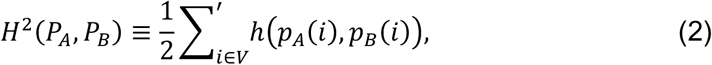

where we use a primed sum notation to indicate that the summation is to be taken as explained above. When applying the above in our calculations, we have to do that separately for the out-neighborhoods and the in-neighborhoods of the nodes, i.e., working with 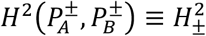.

### Sample complexity

Is defined as the sufficient number of samples needed to correctly differentiate two distributions with a classification error rate *δ* ≤ 0.1 (see next), in a simple binary hypothesis test (Pensia et al., 2022). Suppose *n* samples are drawn from one of two probability distributions *P* and *Q* over support *V*. The test (function) 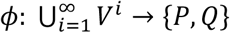 that classifies samples *X* = (*x*^1^, …, *x*^*n*^) ∈ *V*^*n*^ to *P* or *Q*, solves the simple binary hypothesis test problem with sample complexity 𝒩^δ^ (*P, Q*), if it is the smallest number that satisfies

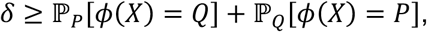

where ℙ_*P*_[*ϕ*(*X*) = *Q*] represents the (failure) probability of classifying the samples as coming from *Q* when drawn from *P*, and vice versa, for the second term (Pensia et al., 2022). Moreover, it is known (Canonne et al., 2019) that it scales as

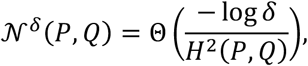

where Θ(⋅) is the Big-Theta notation for asymptotic tight (upper and lower) bound (Cormen, 2007).

Finally, if *ϕ* is the optimal likelihood test and 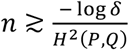 for *δ* ≤ 0.1, the failure probability of the test *ϕ* with *n* samples is less than *δ* (Pensia et al., 2024). In the case of the Scheffé’s test 𝕀(*x*) = 1 if *P*(*x*) ≥ *Q*(*x*) and 0 otherwise (Scheffé, 1956), it is at most 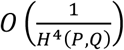 (Pensia et al., 2022, 2024) for *δ* = 0.1, where *O*(⋅) stands for the Big-O notation for the asymptotic upper bound (Cormen, 2007); but it has been found that it can be a tight bound for certain distributions (Suresh, 2020).

### *α*-Rényi divergence

For two probability distributions *P*_*A*_ and *P*_*B*_, the *α*-Rényi divergence (Renyi, 1961) is

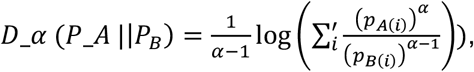

where we already used the primed sum as defined for the Hellinger distance. This function has several properties. In particular, for any *P* and *Q, D*_*α* (*P*||*Q*) increases as a function of *α*. For *α* = 1, *D*_1 (*P*||*Q*) is the Kullback-Leibler divergence, *D*_*KL* (*P*||*Q*). For our purposes, the Rényi divergence of interest is the one with order parameter *α* = 1/2, i.e.,

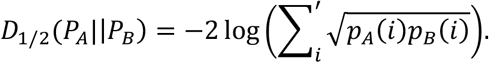

We have chosen *α* = 1/2 because it is the only Rényi divergence that is symmetric. This property is important since in our case, probability mass functions are associated with node neighborhoods. By comparing them, we do not aim to measure their information compression properties – a classical application of information divergence – but to measure how different they are. Moreover, since the probabilities of the same event from both probability mass functions, *p*_*A*_(*i*) and *p*_*B*_(*i*), are multiplied and not divided as in the Kullback-Leibler divergence (or any *α* ≥ 1), we can still measure their dissimilarity even if there are mutually exclusive events, i.e., events that have zero probability in one and non-zero in the other distribution. These two properties make the ½-Rényi divergence useful, since it makes it possible to relate network related quantities such as the similarity between node neighborhoods to information theory and statistical inference. It is associated with the squared Hellinger distance (Nielsen & Boltz, 2011) via equation (1). One simple interpretation of the ½-Rényi divergence *D*_½_ (*P* ║ *Q*) is that is quantifies the error incurred when replacing the distribution *P* with distribution *Q*, assuming that the true distribution is *P* and has many applications. For example, consider two sensors with the same sampling rate, monitoring the occurrence of some phenomenon in a noisy environment. *D*_½_ in this case provides a lower bound on the worst-case misdetection rate of the phenomenon (Shayevitz, 2011). These functions help us to decode the functional organization of the network from an information theoretic perspective (Canonne et al., 2019; Shayevitz, 2011).

### Similarity between directed links

The similarity of out-links *e*_*ij*_ and *e*_*ik*_ from common a node *i* is defined as 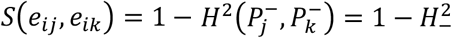. For in-links *e*_*ij*_ and *e*_*ik*_, the similarity measure is 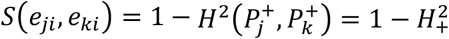. The reason for using the in-neighborhoods of nodes *j* and *k* when measuring the similarity between the out-links *e*_*ij*_ and *e*_*ik*_ from node *i*, is because *e*_*ij*_ and *e*_*ik*_ are in-links to nodes *j* and *k*, and in this way we are comparing in-links with in-links (one belonging to *j*, the other to *k*). In other words, the dissimilarity between directed links is the squared Hellinger distance of the uncommon nodes.

### Extracting the node-community hierarchy from the link-community hierarchy

As stated in the Results section, to derive the node-community hierarchy using the link community algorithm requires specifying how node communities merge through the link merging process. We use two criteria. *i*) The *recurrence criterion* where two node communities merge if they are connected by a link community with recurrent connections, i.e., links (from the same link community) directed both ways between the node communities; and *ii*) the *end-step criterion*, in which node communities that did not merge by the end of the link merging process are then merged pairwise and sequentially. An illustration of the node merging process can be seen in **Figure 4b,c**. Note, the node-community hierarchy depends on these criteria. We choose the recurrence criterion because this is a key property in several multi-agent systems as it allows coordination of information/balance; an example is the mutual inhibition of the neurons in the ventrolateral preoptic and monoaminergic subcortical structures during sleep-wake transitions (Saper et al., 2005). Moreover, it provides the advantage of stability (once two communities are merged, they will not split at any higher threshold value). Therefore, the node-community hierarchy is well-defined.

### Loop entropy

We introduce here the notion of loop entropy, that will provide us with a criterion that allows us to select an “optimal” level in the link-community hierarchy. We start with the definition that trees (which are connected networks whose number of links *M* are related to its number of nodes *N* by the equation *M* = *N* − 1), have no (i.e., zero) community structure. To further simplify matters we will ignore the directionality of the links in what follows. We define the number of excess links *M*_*exc*_ of a network *G* with *M* links and *N* nodes as *M*_*exc*_(*G*) = *M* − *N* + 1 (which is zero for a tree).

Suppose a network *G* is partitioned in *L* mutually exclusive subgraphs 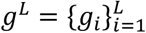 i.e., 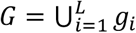 and ∅ = *g*_*i*_ ∩ *g*_*j*_ ∀*i, j* ∈ {1, …, *L*}. Then, the loop entropy, *S*_*L*_, of that partition is defined as

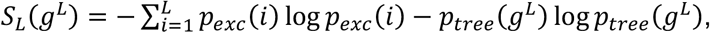

where 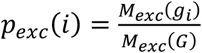 is the probability of sampling excess links in the subgraph *g*_*i*_ from the total number of excess links in *G*. Finally, 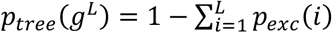 is a normalization factor.

The loop entropy is a measure of the distribution of excess links among the subnetworks, i.e., how much the partition differs from the tree (null) model which does not encode community information (recall, communities must have recurrent links). Since it is a Shannon-Gibbs type entropy, it is maximized when the distribution of excess links of the partition 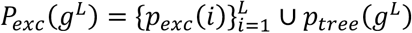 is uniform of the community structure.

Note, we have not proved that *S*_*L*_ can always be computed since it might occur that there are partitions where *M*(*g*) > *M*_*exc*_ (*G*) for a particular subgraph *g*, or that 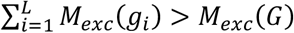 (in this case we no longer work with probabilities). However, we found that the score did not have such issues in the networks analyzed by us. Here we use the loop entropy as a community quality score for link community partitions in various network models and benchmarks (Lancichinetti & Fortunato, 2009) and in all cases the *S*_*L*_ was well defined. Until there is a mathematical proof that *S*_*L*_ can be computed for any arbitrary mutually exclusive partition, we recommend monitoring the *S*_*L*_ values to find possible anomalies, when using it with the link community algorithm. See Supplementary Materials Figure S15 for an example.

### Software packages

We used Python 3.9.18 and libraries SciKit-Learn version 1.1.1, SciPy version 1.13.1, NetworkX version 2.8.8, and Pybind11 2.10.0. For plotting, we used Matplotlib version 3.5.2 and Seaborn version 0.13.2, and R-library ggtree version 3.8.2 (Yu et al., 2017). Benchmarks were generated using the Lancichinetti-Fortunato model for directed, weighted, networks with nodes with multiple communities (Lancichinetti & Fortunato, 2009). We also used the hclust-cpp library that implements the hierarchical clustering algorithm (*HS Niederrhein - Dalitz*, 2020; Müllner, 2013). To compute the omega index, we used a publicly available code (Murray et al., 2012).

## Data availability

The network connectivity data is available from https://core-nets.org/.

## Code availability

The codes used in this study are available and freely accessible on GitHub.

## Acknowledgments

This work was supported by the National Science Foundation (NSF) grant IIS-1724297 (J.M.A., Z.T.), French National Research Agency (ANR) grants A2P2MC ANR-17-NEUC-0004 (K.K., H.K.), ANR-17-FLAG-ERA-HBP-CORTICITY (to H.K.), ANR-19-CE37– 0025-DUAL_STREAMS (to K.K.).

## Author contributions

The authors would like to express their thanks for discussions with B. Molnár, L. Magrou, Y. Hou. J.M.A. and Z.T. designed the research, J.M.A. wrote all the algorithms and ran the simulations, JMA performed the computational and data analysis with guidance from Z.T., K.K. and H.K. provided the experimental datasets, J.M.A. and Z.T. wrote the paper with assistance from K.K. and H.K., and all authors contributed to editing the paper.

## Competing interests

The authors declare no competing interests.

## Additional information

**Extended data** are found below.

## Notes

### Competing Interest Statement

The authors have declared no competing interest.

### Summary of Updates

This revised version of the manuscript introduces several substantial conceptual, methodological, and structural updates. First, the title has been changed from - A principled approach to community detection in interareal cortical networks - to - An information theoretic approach to community detection in dense cortical networks reveals a nested hierarchy - to better reflect the core methodological framework and the main findings of the study. Second, the manuscript has been significantly expanded with a detailed Supplementary Material file. This includes systematic benchmarking of the proposed method against established community detection approaches, as well as additional analyses exploring alternative similarity and distance measures beyond the Hellinger distance. These additions strengthen the validation of the method and clarify its advantages in the context of dense, weighted, and directed cortical networks . Third, the presentation of results has been reorganized. In particular, the original Figure 4 has been revised and split into two separate figures (Figures 4 and 5) to improve clarity and better distinguish between link-community and node-community hierarchical structures. Fourth, the comparison with the SLN-based hierarchy has been removed in this version to maintain a clearer focus on the methodological contributions and the information theoretic framework. While this comparison yielded promising insights into cortical organization, it is beyond the scope of the present manuscript and will be addressed in future work. Finally, several minor revisions have been made throughout the manuscript, including improvements in figure layout, terminology, and overall exposition. These changes are primarily stylistic and do not alter the main results or conclusions. Overall, the revised manuscript provides a clearer, more focused, and more thoroughly validated presentation of the proposed framework for detecting hierarchical community structure in dense cortical networks.

