## Supplementary Materials for "An information theoretic approach to community detection in dense cortical networks reveals a nested hierarchy"

**Abstract**

, This document provides supplementary analyzes supporting the main manuscript submitted to *Network Neuroscience*.

**S4 Supplementary Figures** 18

**S5 Additional References** 29

### S1 Benchmarking

Since there isn't a unique, principled theory for community detection, uncovering the communities of a network relies on various definitions for the notion of a community. Moreover, most methods are heuristic; and while they often share common features even in their output, they typically yield different results. Ultimately, in the absence of a ground truth (when that is the case), they only provide a best estimate decomposition of a system, from the point of view of the specified criteria.

In this section, we present the results of using traditional community detection methods on the edge-complete macaque FLN dataset comprising 40 cortical areas. The algorithms used are: (*i*) the modularity maximization implementation from [5], (*ii*) Infomap [12], and (*iii*) the weighted stochastic block model (WSBM) [2].

At the end of this section, we provide a broad comparison between the results of these methods and those of our own approach (see Table S1).

#### S1.1 Modularity maximization

As part of validating and benchmarking our approach, we first compute community partitions using modularity optimization, a baseline method widely used in community detection. Specifically, we follow the procedure of [5], which applied the modularity maximization method from the Brain Connectivity Toolbox (BCT) [13] to analyze the community structure of the anterograde mouse connectome derived by the Allen Institute, a data set structure comparable to the macaque FLN network.

In this approach, the Louvain algorithm [3] is used to optimize the Reichardt–Bornholdt modularity function [11], while varying the resolution parameter $\gamma$ from low values (coarse partitions) to high values (fine partitions). In our implementation, we sweep $\gamma \in [0, 2.5]$ in increments of 0.1. The directed and weighted Reichardt–Bornholdt modularity is defined as

$$Q(\mathbf{g}, \gamma) = \frac{1}{s} \sum_{i \neq j} \left[ w_{ij} - \gamma \frac{w_i^+ w_j^-}{s} \right] \delta_{g_i g_j}, \tag{S1.1}$$

where $s = \sum_i \sum_j w_{ij}$ is the total edge weight, $\mathbf{g}$ denotes the vector of community labels, $g_i$ is the community assignment of node $i$, $w_{ij}$ is the weighted adjacency matrix entry between source node $i$ and target node $j$, $w_i^+ = \sum_j w_{ij}$ and $w_i^- = \sum_j w_{ji}$ are the sum of out-weights and in-weights of node $i$, respectively, and $\delta$ is the Kronecker delta. The term $w_i^+ w_j^- / s$ is the expected weight of edges from $i$ to $j$ under the directed, weighted configuration model [13].

The Louvain method optimizes the modularity function using a greedy assignment strategy and, in principle, can be applied to both directed and undirected formulations of modularity. In the Brain Connectivity Toolbox, however, the implementation assumes a symmetric modularity matrix. Consequently, for directed networks the modularity contributions are symmetrized prior to optimization, i.e.,

$$Q_{ij} \leftarrow \frac{Q_{ij} + Q_{ji}}{2}, \quad Q_{ji} \leftarrow Q_{ij},$$

where

$$Q_{ij} = \frac{1}{s} \left[ w_{ij} - \gamma \frac{w_i^+ w_j^-}{s} \right] \delta_{g_i g_j}.$$

This step preserves internal consistency of the BCT routine but eliminates edge directionality. As we show below, this restriction leads to meaningful differences between the partitions obtained by modularity maximization and those identified by our method.

For each value of $\gamma$, we perform 1000 independent Louvain runs with randomized node orderings. From these partitions, an affinity matrix is constructed following [5]: the entry for nodes $i$ and $j$ is the fraction of runs in which they co-occur in the same community, weighted by the modularity of each partition and normalized by the total number of runs. This yields an affinity score between node pairs. The consensus partition for that $\gamma$ is then obtained by reapplying the Louvain method 100 times to the affinity matrix and selecting the partition with maximum modularity under the constraint that all affinity scores within a module are at least 0.5.

To assess statistical significance, the same procedure is applied to ensembles of directed configuration model networks that preserve the in- and out-degree sequence of FLN. Following [5], the empirical consensus partition finally retained is the one for which the difference in average modularity between the empirical network and the null ensemble is largest and positive.

**Results** Figure S1 summarizes the results. Panel S1**a** shows the average modularity as a function of $\gamma$ for the macaque FLN network (blue) and the directed configuration model ensemble (red), together with their difference (yellow). The $\gamma$ value that maximizes this difference is indicated by a black dot. This optimal resolution lies slightly below one, indicating that partitions at coarser resolution are most distinct from the null model. The corresponding consensus partition is shown in panel S1**b**.

As mentioned earlier, an important feature of the Brain Connectivity Toolbox implementation is that the modularity matrix is symmetrized prior to optimization, which removes edge directionality. In directed networks such as FLN, where anatomical projections can be highly asymmetric in terms of strength, this symmetrization constitutes a substantive limitation.

For comparability, we also applied our algorithm to a symmetrized version of the edge-complete FLN adjacency matrix. This produced results broadly consistent with modularity maximization (Figure S1**c**). However, a comparison between Figures S1**b** and 5**b** (see main manuscript) shows that the green community in panel S1**b** is subdivided by our method into two sub-communities: one comprising of somatosensory–premotor areas and the other of premotor–motor areas. Such an internal subdivision within the somatosensory, premotor, and motor system has been reported previously [10], but it is not resolved using modularity maximization. We hypothesize that this difference reflects our algorithm's explicit treatment of directional connectivity profiles, which the symmetrization obscures. This illustrates how incorporating directionality can reveal emergent organizational features of cortical networks that would otherwise remain hidden under undirected formulations.

Note also that the partition in Figure S1**c**, obtained at the Goldilocks level as defined by the maximum of loop entropy, largely overlaps with the modularity maximization result (Figure S1**b**). Reaching this partition with modularity maximization requires multiple runs across resolution values, comparisons with null models, and a consensus step. By contrast, our approach arrives at a comparable partition in a single, parameter-free procedure, supporting the observation that the optimal resolution identified by modularity maximization might, in fact, be a good definition for the Goldilocks level.

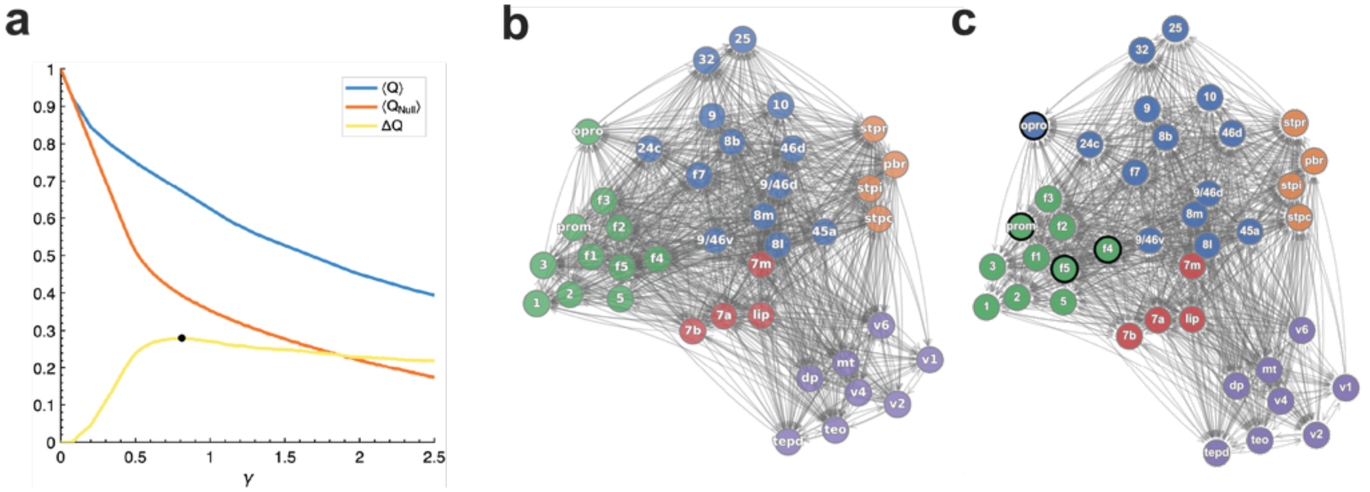

Figure S1: **Results of modularity maximization on the FLN dataset.** (**a**) Average modularity as a function of the resolution parameter $\gamma$ for the macaque FLN network (blue) and directed configuration model ensemble (orange), with their difference shown in yellow. The black dot marks the $\gamma$ value with the largest positive difference, corresponding to the most significant resolution. (**b**) Consensus partition at the optimal $\gamma$ value, obtained from the Brain Connectivity Toolbox implementation of the Louvain method, which symmetrizes the modularity matrix prior to optimization. (**c**) Partition obtained by applying our algorithm to a symmetrized version of the FLN adjacency matrix. While broadly consistent with panel **b**, when applied to the original directed FLN matrix, our method subdivides the green community into somatosensory–premotor and premotor–motor subcommunities, highlighting the role of directional connectivity profiles, whereas the other method does not.

#### S1.2 Infomap

Infomap [12] is a community detection method based on information theory. It identifies modules by minimizing the description length of a random walker navigating the network. Connection weights are normalized into transition probabilities, and the optimal partition is the one that yields the most compressed description of the walker's trajectory. The description length $L(M)$ is defined by the *map equation* (Eq. S1.2):

$$L(M) = q_{\curvearrowright} H(\mathcal{Q}) + \sum_{i=1}^{m} p^i_{\circlearrowright} H(\mathcal{P}^i), \tag{S1.2}$$

where $q_{\curvearrowright}$ is the probability of exiting a module, $H(\mathcal{Q})$ is the Shannon entropy of inter-module transitions, $p^i_{\circlearrowright}$ is the probability of being within module $i$, and $H(\mathcal{P}^i)$ is the entropy of transitions inside module $i$. The optimal partition $M$ minimizes $L(M)$.

Because an exhaustive search over all possible partitions is computationally intractable, Infomap relies on a greedy, agglomerative heuristic: modules are iteratively merged whenever this reduces the value of the map equation. In contrast to the BCT implementation of modularity maximization, Infomap natively supports directed and weighted networks, since transition probabilities in the underlying random walk incorporate both edge direction and weight.

Infomap controls the granularity of communities through the Markov time parameter, which dictates how long a random walker explores the network. Shorter times yield fine-grained partitions, whereas longer times produce coarser ones. We varied the parameter from 0.2 to 3 in steps of 0.1 (Fig. S2**a**). For each value, we ran the algorithm 100 times and, as in the maximum modularity case, constructed a weighted co-classification matrix based on description length, normalized by the total description length across all instances. Because Infomap seeks partitions with minimum description length, we inverted the non-zero entries of the co-classification matrix and derived the final partition using the Louvain algorithm. Across Markov times, we also computed the average description length from configuration-model randomized networks, using the same number of runs as for the macaque FLN network. Finally, we selected the partition with the largest positive difference between the description length in the data and the null model, i.e., the level at which the empirical network can be compressed more efficiently than the configuration model. In this case, the maximum difference occurred at Markov time 0.9.

**Results** The resulting partition is shown in Figure S2**b**, where Infomap identifies seven modules. Within the prefrontal–cingulate region, the main difference, relative to ours, concerns the assignment of three areas: area 10 is grouped with areas 25–32 in our hierarchy, whereas areas 46d and 9/46d are clustered with 8m–8L–9/46v–45a. Given that this region comprises 13 areas in total, the discrepancy is minor, and both methods identify three subcommunities. Similarly to modularity maximization, Infomap does not separate the somatosensory–premotor and premotor–motor systems. These subdivisions are supported by anatomical and functional evidence, suggesting that different optimization principles may emphasize distinct organizational features of the network. More generally, Infomap and Louvain provide effective heuristic partitions, whereas our approach is based on a probabilistic interpretation of connectivity profiles, offering a complementary perspective on cortical network organization.

#### S1.3 Weighted stochastic block model

The weighted stochastic block model (WSBM) [2] is a generative approach to community detection. It assumes that both edge existence and edge weights are generated from probability distributions whose parameters depend only on the community memberships of the nodes at their endpoints. In this way, the WSBM extends the classical stochastic block model, which accounts only for binary edges, to networks with weighted connections.

Let $A$ denote the adjacency matrix, $z = \{z_i\}$ the latent community assignments of the nodes, and $\theta$ the parameters of the block–block edge weight distributions. Inference proceeds by approximating the true posterior $p(z, \theta \mid A)$ with a tractable distribution $q(z, \theta)$. The optimization problem can be written as minimizing the Kullback–Leibler (KL) divergence between these two distributions:

$$\mathcal{L}(q) = \mathrm{KL}(q(z, \theta) \parallel p(z, \theta \mid A)) \, . \tag{S1.3}$$

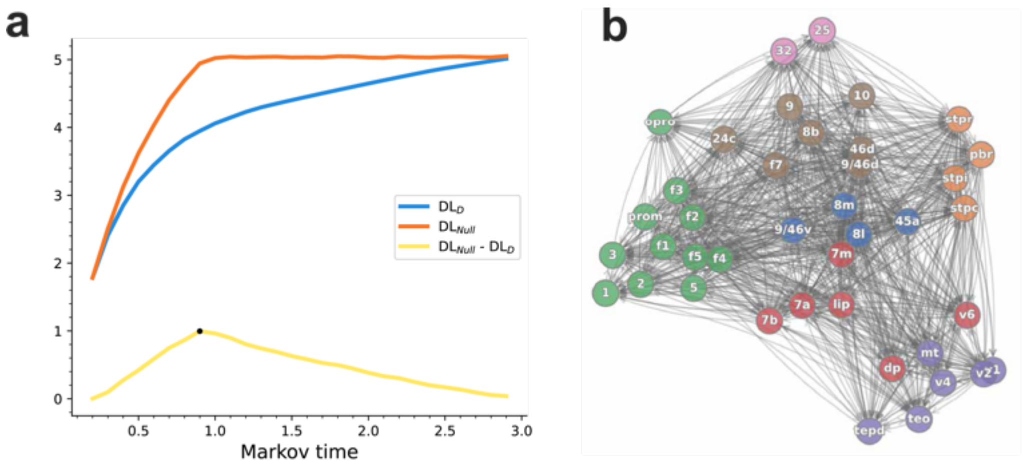

Figure S2: **Infomap results on the FLN dataset.** **a**, Description length as a function of Markov time for the data (blue), the configuration-model null (orange), and their difference (yellow). The maximum difference is marked with a dot. **b**, The selected partition contains seven modules. Differences from our method are mainly in three prefrontal areas (10, 46d, 9/46d) and in the assignment of cingulate areas 25 and 32. As with modularity maximization, Infomap does not resolve the subdivision between somatosensory–premotor and premotor–motor systems. This illustrates both the strength of Infomap in handling directed and weighted edges and its limitation in capturing biologically meaningful subdivisions.

Here, $p(A, z, \theta) = p(A \mid z, \theta)\, p(z)\, p(\theta)$, where $p(A \mid z, \theta)$ is defined by the chosen family of edge-weight distributions, and $p(z)$ and $p(\theta)$ are independent priors on the community assignments and block parameters, respectively. The expectation is taken with respect to the variational distribution $q(z, \theta)$.

The marginal log-likelihood of the network,

$$\log p(A) = \int dq(z, \theta)\, \log p(A),$$

can be decomposed into two terms:

$$\log p(A) = \mathcal{G}(q) + \mathcal{L}(q),$$

where

$$\mathcal{G}(q) = \mathbb{E}_q[\log p(A \mid z, \theta)] + \mathbb{E}_q\left[\log \frac{p(z)p(\theta)}{q(z,\theta)}\right].$$

Since $\log p(A)$ is independent of $q$, maximizing $\mathcal{G}(q)$ is equivalent to minimizing $\mathcal{L}(q)$. In practice, inference therefore reduces to maximizing $\mathcal{G}(q)$, which is computationally tractable.

A key feature of the WSBM is the inclusion of a mixing parameter $\alpha \in [0, 1]$, which balances the contribution of two components in the likelihood: one distribution governing edge existence and another governing edge weights. The edge-existence distribution is often taken to be Bernoulli for binary presence or absence of edges, but in principle other exponential-family distributions (e.g., normal, Poisson) can also be used. When $\alpha = 0$, the model reduces to the classical (unweighted) stochastic block model. When $\alpha = 1$, the model considers only edge weights, ignoring edge presence. Intermediate values of $\alpha$ combine both sources of information, allowing the WSBM to flexibly model weighted networks with heterogeneous sparsity patterns.

By varying the number of communities (blocks), the WSBM jointly optimizes the assignment vector $z$, the distribution parameters $\theta$, and the mixing parameter $\alpha$, selecting the partition that best explains the observed edge structure and weights under this probabilistic framework.

**Results** In our implementation, we varied the number of communities $R$ from 2 to 10 and ran the WSBM algorithm 100 times for $\alpha = 0$ and $\alpha = 1$, corresponding to the two limiting cases that use only edge statistics or only weight statistics, respectively. We modeled edge existence with a Bernoulli distribution and edge weights with a Lognormal distribution. For each $R$, we selected the trial with the highest value of $\mathcal{G}$ (Figure S3**a**). The partitions maximizing $\mathcal{G}$ correspond to $R = 5$ for $\alpha = 0$ and $R = 2$ for $\alpha = 1$.

Figures S3**b**–**c** show the resulting partitions for each $\alpha$ value. For $\alpha = 0$, which relies only on edge statistics, the five-community partition is difficult to interpret and diverges from both our results and those obtained with modularity maximization and Infomap. A notable similarity is the green community, grouping somatosensory–premotor–motor areas, consistent with the other methods. However, the red community combines superior temporal polysensory regions (e.g., STPr, PBr) with visual areas (e.g., MT, DP, TEO), which seems unlikely to represent a functional system given the large spatial separation of its components. Similar concerns apply to the orange and blue communities, which span widely separated regions across premotor, parietal, prefrontal, and superior temporal polysensory cortex.

For $\alpha = 1$ (Figure S3**c**), the two-community partition captures a coarse core–periphery organization: one group contains visual, somatosensory, and cingulate areas, while the other comprises prefrontal, premotor, motor, and superior temporal polysensory regions. This split broadly reflects a division between low-level and high-level cortical areas. While plausible, it oversimplifies the network's structure, as visual and somatosensory regions are engaged in orthogonal functions (encoding visual versus tactile information) and should not be grouped into a single functional subsystem.

Overall, we find that WSBM produces partitions that differ from those obtained with modularity maximization, Infomap, and our approach, often yielding communities that are less spatially or functionally localized. In anatomical neural networks, both edge presence and connection weights carry important information for defining communities, which makes the choice of the mixing parameter $\alpha$ non-trivial. These observations suggest that, while WSBM provides a flexible generative framework, its outcomes in this context emphasize different organizational features and may be less aligned with known spatial and functional subdivisions of the macaque cortex.

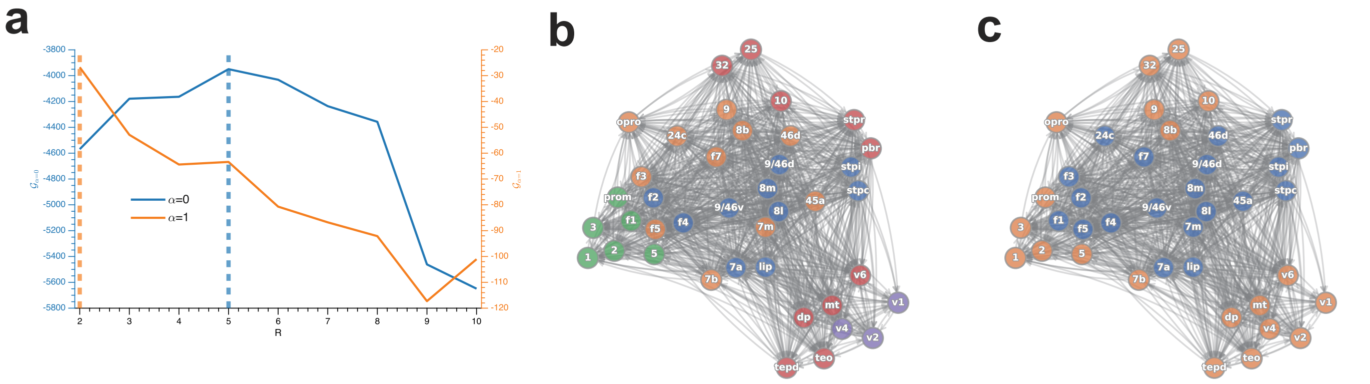

Figure S3: **Results of the weighted stochastic block model (WSBM) applied to the FLN network.** (**a**) Maximum value of $\mathcal{G}$ across 100 runs for each number of communities $R$ (2–10) and for $\alpha = 0$ (edges only) and $\alpha = 1$ (weights only). The optimal partitions correspond to $R = 5$ for $\alpha = 0$ and $R = 2$ for $\alpha = 1$. (**b**) Partition with maximum $\mathcal{G}$ for $\alpha = 0$, yielding five communities. A somatosensory–premotor–motor cluster (green) aligns with other methods, but other modules—for example, the red community linking superior temporal polysensory and visual areas—are difficult to interpret functionally. (**c**) Partition with maximum $\mathcal{G}$ for $\alpha = 1$, producing a two-community structure that reflects a coarse division between low-level (visual, somatosensory, cingulate) and high-level (prefrontal, premotor, motor, superior temporal polysensory) areas. While this captures a core–periphery split, it oversimplifies cortical organization by grouping functionally distinct systems.

#### S1.4 Comparison of methods

Having presented the results of modularity maximization, Infomap, WSBM, and our algorithm, we now summarize their main similarities and differences (see Table S1 for a side-by-side comparison).

Despite relying on different optimization principles, both modularity maximization and Infomap identified broadly consistent communities, capturing spatially and functionally coherent clusters such as a somatosensory–premotor–motor module and a set of visual areas. Our method produced comparable partitions and, by explicitly incorporating directionality, further subdivided these clusters into somatosensory–premotor and premotor–motor regions. This highlights how different approaches emphasize different organizational features of the same dataset.

In contrast, WSBM generated results that diverged more strongly from the other methods. For $\alpha = 0$ (edge statistics only), the five-community solution included clusters that were spatially dispersed and less interpretable (e.g., grouping superior temporal polysensory and visual regions). For $\alpha = 1$ (weight statistics only), the two-community solution revealed a coarse split between low-level and high-level areas, but merged functionally distinct systems. These outcomes illustrate the sensitivity of generative approaches to parameter choices, and show that they may reveal coarse-grained organization while not always recovering spatially localized modules.

Taken together, these benchmarks show that different community detection algorithms, by design, highlight distinct facets of cortical network organization: modularity maximization and Infomap capture broad-scale modules; WSBM emphasizes generative block-structure assumptions; and our method leverages connectivity profile similarity and directionality to reveal a nested hierarchy of modules. Importantly, in the absence of ground-truth partitions of cortical areas, none of these outcomes should be viewed as definitive. Instead, they provide complementary hypotheses about brain organization that can be evaluated and refined by future multimodal evidence.

Finally, our method's distinctive contribution is to connect anatomical connectivity with a metric-based hierarchical framework: the squared Hellinger distance between node connectivity profiles defines a natural measure of similarity, embedding the network into a metric space with an associated depth scale. This complements approaches based on global cost-function optimization, such as modularity maximization, Infomap, and WSBM, by providing an alternative, information-theoretic perspective on multiscale cortical organization.

Table S1: Comparison of community detection methods applied to the FLN network. Each method emphasizes different aspects of cortical organization, reflecting its underlying principles.

| **Method** | **Principle** | **Strengths** | **Limitations / Results in FLN** |
|---|---|---|---|
| Our method | Information-theoretic, feature-based hierarchical clustering incorporating weights and directions | Incorporates edge directionality; produces a nested hierarchy; identifies subdivisions such as premotor–motor vs. somatosensory–premotor | Computationally more demanding; results largely consistent with modularity/Infomap at broad scales, with additional subdivisions reflecting asymmetric connectivity |
| Modularity maximization | Greedy optimization of the Reichardt–Bornholdt modularity via Louvain (BCT implementation) | Widely used baseline; captures coherent large-scale modules such as somatosensory–premotor–motor and visual; applied to multiple connectomes including mouse [5] | In the BCT version, the modularity matrix is symmetrized, discarding edge directionality; most informative partitions obtained for $\gamma < 1$ |
| Infomap | Minimization of the map equation; information-theoretic coding of random walks via description length | Detects flow-based communities; does not require specifying the number of modules; incorporates directionality and weights | Broad agreement with modularity at large scale; in FLN, produced partitions of similar granularity without resolving all observed subdivisions |
| WSBM | Probabilistic block model with edge/weight statistics; parameter $\alpha$ interpolates between them | Flexible generative framework; can capture block-structured and core–periphery organizations | Results depend on parameter choices; in FLN, $\alpha = 0$ yielded 5 communities (some spatially dispersed), while $\alpha = 1$ produced 2 communities reflecting a coarse low- vs. high-level division |

### S2 Similarity Analysis

Our method relies on features extracted from local connectivity profiles among cortical areas. We use the Hellinger distance because it is a proper metric for comparing probability distributions, making it well-suited for FLN inputs. However, its interpretation, particularly its link to the notion of "sample complexity" (see main text), may feel less intuitive than more familiar measures such as Jaccard indices, cosine similarity, or correlations. To place our results in context, we therefore applied our algorithm with alternative similarity measures: weighted Jaccard, Jaccard probability, cosine similarity, and the Pearson correlation. Unless otherwise specified, similarities between feature vectors were computed using the primed-sum convention of Equation 2 in the main text. A side-by-side comparison of the measures is provided in Table S2.

#### S2.1 Jaccard index

The *Jaccard index* ($J_{\mathcal{S}}$) measures the overlap between two sets:

$$J_{\mathcal{S}}(A,B) = \frac{|A \cap B|}{|A \cup B|}, \qquad J_{\mathcal{S}} \in [0,1],$$

with the associated distance $d_J = 1 - J_{\mathcal{S}}$. For binary feature vectors $\mathbf{A}, \mathbf{B} \in \{0,1\}^N$, the index can be written as

$$J_{\mathcal{S}}(\mathbf{A},\mathbf{B}) = \frac{\mathbf{A} \cdot \mathbf{B}}{\|\mathbf{A}\|_2^2 + \|\mathbf{B}\|_2^2 - \mathbf{A} \cdot \mathbf{B}}. \tag{S2.1}$$

Because $J_{\mathcal{S}}$ compares sets, it is most suitable for binary (or binarized) data. In FLN, however, *weights* carry essential information about interareal communication, so the plain Jaccard index is not an appropriate measure of similarity between connectivity profiles.

#### S2.2 Weighted Jaccard

To incorporate weights, the *weighted Jaccard* index ($J_{\mathcal{W}}$) adopts a min–max formulation:

$$J_{\mathcal{W}}(\mathbf{A},\mathbf{B}) = \frac{\sum_{i=1}^{\prime N} \min(a_i, b_i)}{\sum_{i=1}^{\prime N} \max(a_i, b_i)},$$

for non-negative vectors $\mathbf{A}, \mathbf{B} \in \mathbb{R}^N_{\geq 0}$ with $\sum_i a_i > 0$ and $\sum_i b_i > 0$. If $a_i, b_i \in \{0,1\}$ for all $i$, then $J_{\mathcal{W}} = J_{\mathcal{S}}$. As with $J_{\mathcal{S}}$, we have $J_{\mathcal{W}} \in [0,1]$, and the associated distance $d_{J_{\mathcal{W}}} = 1 - J_{\mathcal{W}}$ defines a metric.

**Results** At the Goldilocks level (maximum loop entropy), the partition induced by $J_{\mathcal{W}}$ (Figure S4**a**) shares broad features with those obtained using $H^2$ and Infomap, including a large prefrontal–cingulate module and a separation between somatosensory–premotor–motor and visual regions. Some differences appear in the details: (*i*) $J_{\mathcal{W}}$ groups area F1 with areas 3 and 2 (primary somatosensory), whereas it is well-known that F1 is functionally distinct; (*ii*) $H^2$ and external evidence indicate that area 1 has a mixed somatosensory–motor role [10], while in $J_{\mathcal{W}}$ it is not grouped with the other primary somatosensory areas, only premotor areas; and (*iii*) $J_{\mathcal{W}}$ places V6 and DP with parietal areas (7a/7m), a pattern also found with Infomap. These similarities and differences suggest a connection between the random-walk/description-length principle used by Infomap and the min–max overlap emphasized by $J_{\mathcal{W}}$, with each measure highlighting distinct aspects of the underlying connectivity profiles.

A practical caveat is that $J_{\mathcal{W}}$ is biased when feature vectors represent probability mass functions: it can underestimate overlap relative to the set-based Jaccard [8]. For instance, if $\mathbf{A}$ and $\mathbf{B}$ are normalized by their numbers of nonzeros and $|\mathbf{A}| \neq |\mathbf{B}|$, then $J_{\mathcal{W}}(\mathbf{A},\mathbf{B}) < J_{\mathcal{S}}(\mathbf{A},\mathbf{B})$. This motivates the probability-aware variant below.

#### S2.3 Jaccard probability

The *Jaccard probability* ($J_{\mathcal{P}}$) extends Jaccard to probability vectors $\mathbf{A}, \mathbf{B} \in \mathbb{R}^N_{\geq 0}$ with $a_i, b_i \geq 0$ and $\sum_i a_i = \sum_i b_i = 1$:

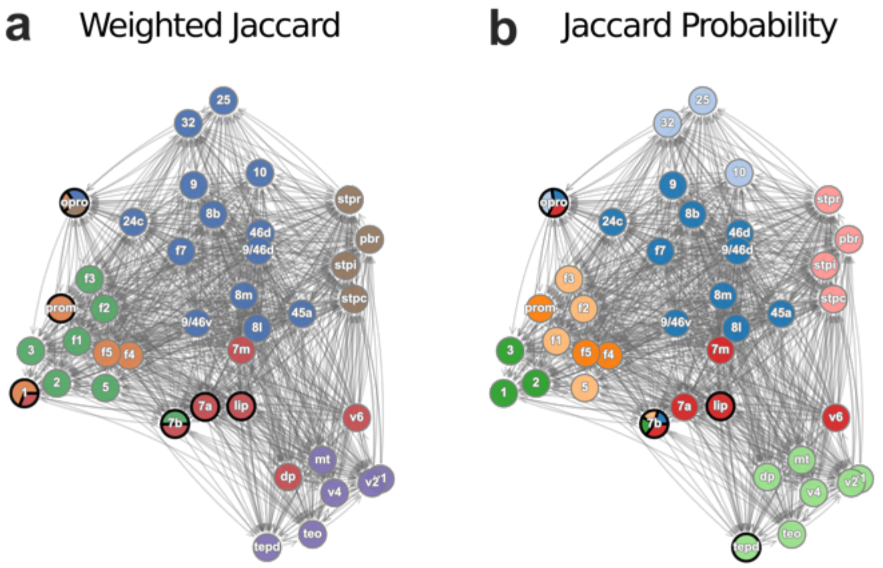

Figure S4: **Goldilocks partitions from Jaccard indices.** Community structure of the FLN network at maximum loop entropy using (**a**) the weighted Jaccard index and (**b**) the Jaccard probability index. Both measures recover broad organizational features also observed with $H^2$ and Infomap, including a large prefrontal–cingulate community and the separation of somatosensory, premotor, and motor systems. Weighted Jaccard groups area F1 with primary somatosensory areas (3 and 2), whereas Jaccard probability assigns area 1 with areas 3 and 2. The similarity between the Jaccard probability and $H^2$ partitions suggests that both measures capture shared structural information embedded in connectivity probability distributions.

$$J_{\mathcal{P}}(\mathbf{A},\mathbf{B}) = {\sum_{\substack{i=1\\ a_i,b_i>0}}^{N}}' \frac{1}{{\sum'}_{j=1}^{N} \max\left(\frac{a_j}{a_i},\, \frac{b_j}{b_i}\right)},$$

which retains desirable properties for probability mass functions and mitigates the underestimation issue of $J_{\mathcal{W}}$ [8]. In our context, applying $J_{\mathcal{P}}$ to FLN profiles preserves their probabilistic interpretation.

**Results** The node-community partition at maximum loop entropy using $J_{\mathcal{P}}$ (Figure S4**b**) closely resembles the $H^2$ partition, while showing fine-grained differences within the somatosensory–premotor–motor and cingulate systems. In agreement with $H^2$, $J_{\mathcal{P}}$ separates a premotor–motor cluster (F1, F2, F3, area 5) from the remaining somatosensory-premotor areas. Unlike $H^2$, it assigns area 1 with areas 3 and 2 (primary somatosensory), without indicating its association to motor regions [10]. As in $H^2$, it also groups V6 with 7a/7m (without DP), unlike $J_{\mathcal{W}}$ and Infomap. These consistencies suggest that both the premotor–motor subdivision and the asymmetric treatment of V6 and DP are characteristic features revealed by probability-preserving similarity measures.

The similarity between $J_{\mathcal{P}}$ and $H^2$ is more apparent in their hierarchical dendrograms (Figure S5): branch arrangements are nearly identical to Figure 5 in the main text, with two differences: (i) area 1 lies closer to areas 2/3 under $J_{\mathcal{P}}$, whereas under $H^2$ it lies near the root beside OPRO and 7b; and (ii) absolute merge heights are larger under $J_{\mathcal{P}}$, indicating greater dissimilarity values in that metric.

Overall, $J_{\mathcal{P}}$ and $H^2$ produce similar hierarchical structure. We prefer the Hellinger distance because it is directly related to the 1/2-Rényi divergence, $D_{1/2}$, which measures distributional differences on a logarithmic scale (akin to KL divergence) and, in our data (Figure 3**b**), scales linearly with interareal distance, an interpretable information-theoretic relationship between profile divergence and spatial separation.

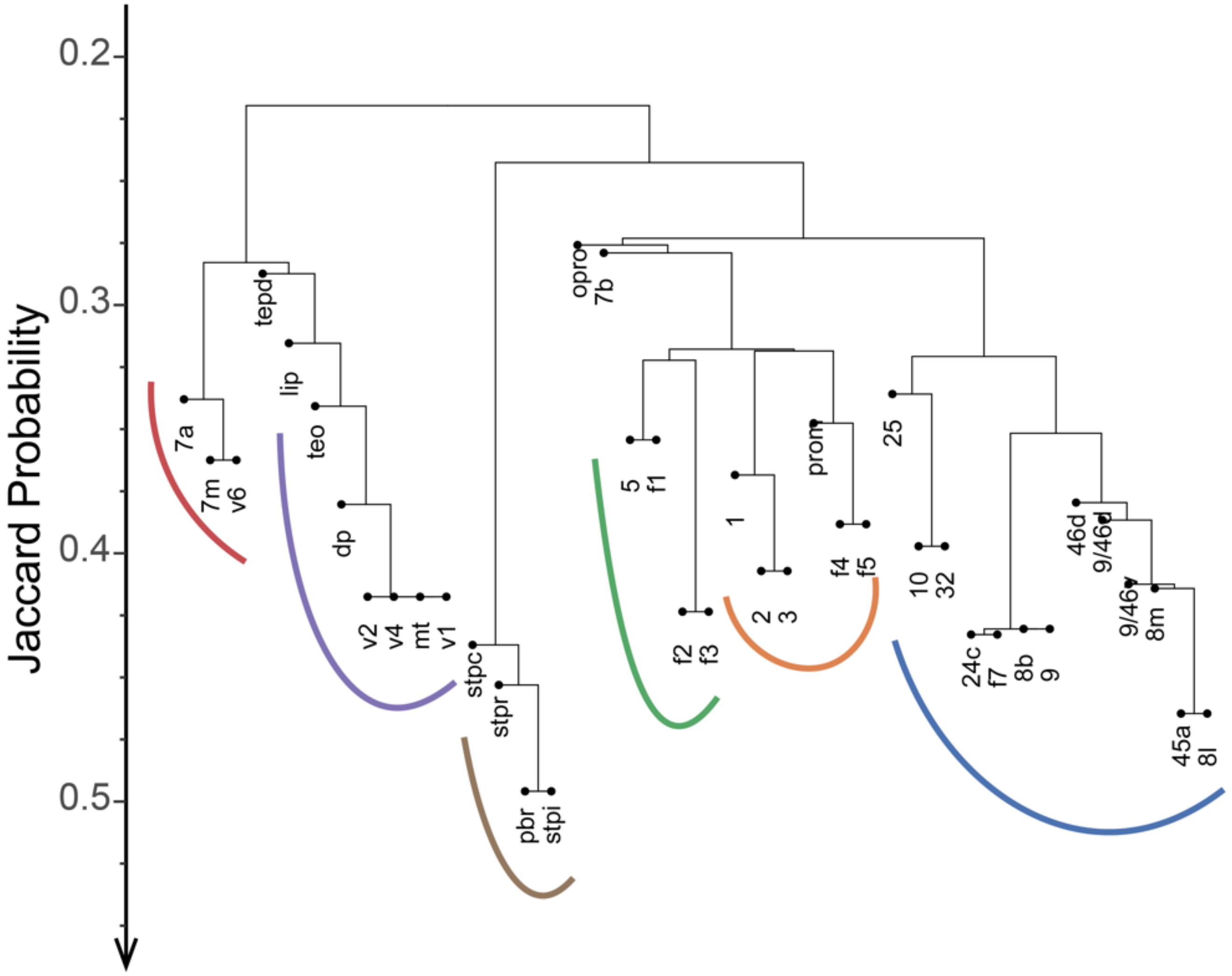

Figure S5: **Hierarchical community structure from the Jaccard probability index.** Dendrogram of the FLN network using the Jaccard probability index. The overall branching pattern closely parallels the $H^2$ hierarchy (Figure 5 in the main text), with both showing six main branches and a comparable order within them. The main differences are: (1) area 1 clusters with areas 3 and 2 under $J_{\mathcal{P}}$, whereas in the $H^2$ dendrogram it is placed closer to OPRO and 7b; and (2) merge heights are generally larger under $J_{\mathcal{P}}$, indicating weaker baseline contrast between community structures compared to $H^2$.

##### S2.4 Cosine similarity and Pearson correlation

Another widely used approach for quantifying vector similarity is cosine similarity, which measures the angular separation between vectors:

$$\cos(\theta_{\mathbf{AB}}) = \frac{\mathbf{A} \cdot \mathbf{B}}{\sqrt{\|\mathbf{A}\|_2^2 \|\mathbf{B}\|_2^2}},$$

where $\mathbf{A}, \mathbf{B} \in \mathbb{R}^N$ and $\cos(\theta) \in [-1, 1]$. Unlike the previous measures, cosine similarity can accommodate feature vectors with negative entries. The Pearson correlation coefficient, $\rho$, can be viewed as cosine similarity computed on $z$-scored vectors:

$$\rho_{\mathbf{AB}} = \cos(\theta_{Z(\mathbf{A}),Z(\mathbf{B})}),$$

where $Z(\mathbf{X}) = (\mathbf{X} - \mu_{\mathbf{X}})/\sigma_{\mathbf{X}}$ is the standardized version of $\mathbf{X}$, with mean $\mu_{\mathbf{X}}$ and standard deviation $\sigma_{\mathbf{X}}$. The range of $\rho$ is also $[-1, 1]$.

**Results** The Goldilocks partitions obtained with cosine similarity and Pearson correlation are shown in Fig. S6**a**–**b**. Both measures produced broadly similar outcomes at the cluster and NOC levels. In agreement with Infomap, they revealed a fine-grained partition of the prefrontal–cingulate cortex, including a central cluster with areas 8L, 8M, 45a, and 9/46v. However, both also yielded configurations that appear less consistent with current anatomical and functional knowledge. For instance, under cosine similarity, areas 7m and V6 were grouped with early visual areas V1, V2, and V4. While 7m and V6 are associated with the dorsal ("where") stream, V1–V4 are crucially involved in early and ventral ("what") stream processing, making this grouping biologically implausible. A related issue appeared with Pearson correlation, where areas 7m, V6, and V4 clustered together despite V4's strong ventral stream role. In addition, both measures placed the primary motor area F1 in the same module as V1 and V2, which are functionally distinct.

Overall, these results suggest that while cosine similarity and Pearson correlation can capture certain structural aspects of connectivity profiles, they may not provide biologically as meaningful partitions of cortical areas at the mesoscopic level.

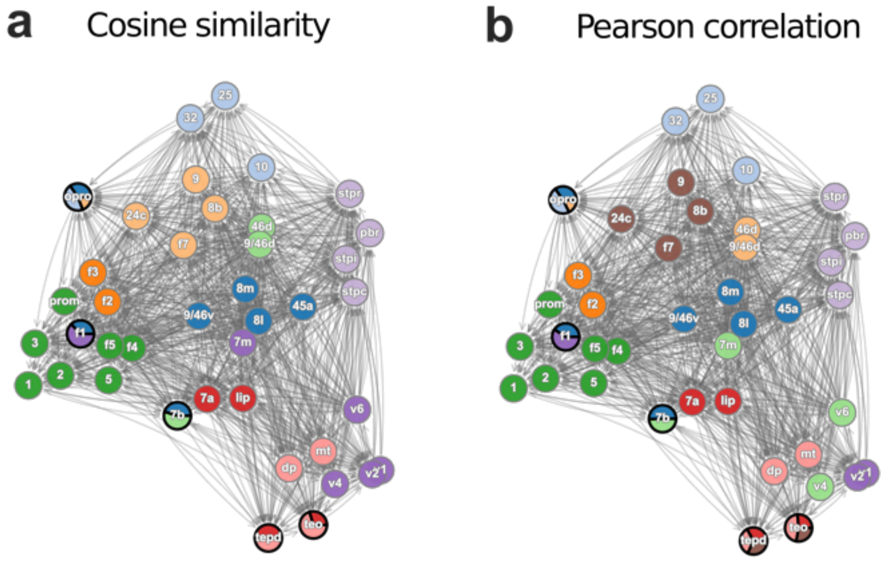

Figure S6: **Goldilocks partitions from cosine similarity and Pearson correlation.** Community structure of the FLN network at maximum loop entropy using (**a**) cosine similarity and (**b**) Pearson correlation. Both measures reveal fine-grained subdivisions in the prefrontal–cingulate cortex, consistent with Infomap. However, they also generate biologically less plausible clusters, such as grouping dorsal-stream areas (7m, V6) with ventral-stream visual areas (V1–V4), or placing the primary motor area F1 with early visual cortex (V1–V2). These results suggest that while cosine- and correlation-based similarities capture certain structural aspects, they do not preserve the probabilistic interpretation of connectivity profiles and may yield less meaningful partitions at the mesoscopic level.

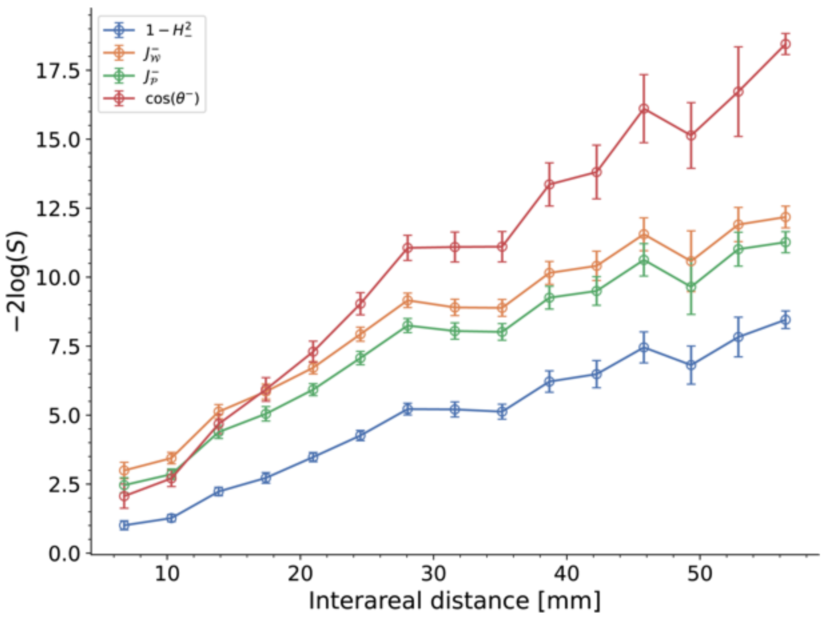

Figure S7: **Scaling of similarity measures with interareal distance.** Average dissimilarity between cortical areas (15-bin averages; error bars = SEM) after applying the 1/2-Rényi transformation, $-2\log(S)$, to different similarity scores: $1 - H^2_-$ (blue), $J^-_\mathcal{W}$ (green), $J^-_\mathcal{P}$ (orange), and $\cos(\theta^-)$ (red). All measures show an approximately linear increase with interareal distance. However, the logarithmic interpretation is strictly valid only for $H^2$, where the curve corresponds directly to the 1/2-Rényi divergence. Notably, $H^2$ yields the lowest average dissimilarity across distances, followed by $J_\mathcal{P}$, while $J_\mathcal{W}$ and cosine similarity assign progressively larger dissimilarities. This pattern suggests that probability-preserving measures ($H^2$ and $J_\mathcal{P}$) evaluate differences more gently and produce community structures that align more closely with known cortical organization.

#### S2.5 Similarity scores scaling

We have examined the results of several similarity scores and analyzed their structural implications. In our framework, we adopted the Hellinger distance as the reference measure because it has a principled connection to the 1/2-Rényi divergence, which quantifies dissimilarity between probability distributions on a logarithmic scale (analogous to the KL divergence). To obtain a global comparison of similarity scores, we applied the 1/2-Rényi transformation (Eq. 1 in the main text), $-2\log(S)$, to in-neighborhoods of the 40 macaque injected areas for each score: $1 - H^2_-$, $J^-_\mathcal{W}$, $J^-_\mathcal{P}$, and $\cos(\theta^-)$. We note that (*i*) in-neighborhoods ($-$) were used because this direction has complete information; (*ii*) the transformation is well-defined because the selected similarity scores all take values in $[0, 1]$ for the non-negative FLN weights; (*iii*) Pearson correlation was excluded since it can yield negative values, which are incompatible with the logarithmic transformation; and (*iv*) the logarithmic interpretation is strictly valid only for $H^2$, while for the other scores it should be regarded as heuristic.

Figure S7 shows the 15-bin averages of $-2\log(S)$ as a function of interareal distance. All similarity scores display an approximately linear increase with distance. Notably, the blue curve (Hellinger distance) indicates the lowest average dissimilarity between cortical profiles, followed by $J_\mathcal{P}$, then $J_\mathcal{W}$, and finally $\cos(\theta)$, which produces the largest dissimilarity values at intermediate and long distances.

These findings help explain why $H^2$ and $J_\mathcal{P}$ produced the most biologically interpretable community structures: both measure similarity in a comparatively "gentle" manner while preserving the probabilistic interpretation of connectivity weights. In particular, $H^2$ reduces the influence of small discrepancies by comparing the square roots of probability distributions, thereby smoothing dissimilarities. This property may be critical in noisy neuroscience datasets, where strict measures can exaggerate small differences, as seen with cosine similarity, where minor fluctuations in feature vectors can strongly affect angular separation.

#### S2.6 Conclusions

Taken together, these analyses show that different similarity measures highlight distinct aspects of the FLN connectivity profiles. The weighted Jaccard, and the Jaccard probability indices recovered several organizational features also detected with $H^2$, including the premotor–motor subdivision and the asymmetric treatment of V6 and DP observed with $H^2$ and $J_{\mathcal{P}}$. Among these, Jaccard probability most closely resembled the hierarchical organization obtained with $H^2$, consistent with both measures preserving the probabilistic interpretation of connection weights. Cosine similarity and Pearson correlation, in turn, emphasized finer subdivisions within the prefrontal and cingulate regions, but also produced groupings that differ from current anatomical and functional expectations, such as dorsal–ventral stream distinctions or the separation of motor and visual systems. This suggests that each measure offers a complementary perspective on the network, with probability-preserving metrics ($H^2$ and $J_{\mathcal{P}}$) providing partitions more aligned with known large-scale cortical organization.

A global comparison of similarity scores further clarifies these differences. Using the 1/2-Rényi transformation $-2\log(S)$, we found that $H^2$ consistently produced the lowest average dissimilarity across interareal distances, followed by $J_{\mathcal{P}}$, then $J_{\mathcal{W}}$, and finally cosine similarity. This ordering reflects how each measure evaluates overlap: $H^2$ (and $J_{\mathcal{P}}$) provide a comparatively gentle assessment of differences by respecting the probabilistic nature of connectivity profiles, whereas weighted Jaccard and cosine similarity penalize mismatches more harshly. In particular, $H^2$ reduces the impact of small discrepancies by comparing square roots of probabilities, a property that may be especially important in noisy neuroscience datasets where strict comparisons can obscure underlying structure. Overall, these comparisons suggest that measures preserving the probabilistic interpretation of FLN weights produce community structures that align more closely with anatomical and functional evidence. This reinforces our choice of the Hellinger distance, which not only shares these desirable properties but also has a principled link to the 1/2-Rényi divergence and reveals a novel linear relationship between connectivity profile divergence and interareal distance.

### S3 Extended Methods

#### S3.1 Link community algorithm example in a toy model

The link-community algorithm works as follows. Initially, each link forms its own community, and the link-community merging threshold value is set to $th = 0$. We define the distance between two link communities $C_i$ and $C_j$, $D(C_i, C_j)$, as the minimum distance over all pairs of links, $u \in C_i$ and the other from $v \in C_j$, i.e,

$$D(C_i, C_j) = \min_{u \in C_i, v \in C_j} d(u, v),$$

where $d(u, v)$ is the distance between links $u$ and $v$ (in terms of either $H^2_+$ or $H^2_-$) . This procedure corresponds to the *single-linkage criterion* in the clustering literature [9]. Next, the threshold $th$ is increased until it reaches the smallest value among all current link-community distances, i.e.,

$$th = \min_{i,j} D(C_i, C_j).$$

At this point, all pairs of communities separated by this distance value $th$ are merged into new communities at the next hierarchical level, and the community list is updated. This process is repeated iteratively until only one community remains.

We illustrate the algorithm on a simple toy model consisting of a directed binary network with three nodes and five links (S8**a**). First, each link is assigned to its own link community. Then, we construct a complete graph whose nodes are the network's links, with edge weights given by the pairwise dissimilarities between them (S8**b**). The algorithm starts by selecting the lowest dissimilarity in this graph as the initial threshold $th$ (i.e., maximum similarity, $th = 0$). Link communities whose dissimilarities equal $th$ are then merged pairwise (link-merging process), forming new communities (S8**b**-1–3). The threshold is then increased to the next minimum link-community dissimilarity (in the example, $th = 0.29$), and the merging process repeats (S8**b**-4). As long as the dissimilarity between two members, one from each link community, equals $th$, the communities are merged. This again corresponds to the single-linkage criterion [9]. The merging process continues until only one link community remains (S8**b**-5). Finally, the information about which link communities merged at which threshold values is used to construct a dendrogram, referred to as the link-community hierarchy (S8**c**).

#### S3.2 An organizational analogy

To illustrate the relationship between the link and node level community hierarchies we consider the university structure in terms of interactions between professors at a university. At the smallest research group level,

Table S2: Comparison of similarity measures applied to FLN connectivity profiles. Each measure emphasizes different aspects of the data, reflecting its mathematical assumptions.

| **Similarity measure** | **Principle** | **Strengths** | **Limitations / Results in FLN** |
|---|---|---|---|
| Hellinger distance ($H^2$) | Information-theoretic metric; compares probability distributions; directly related to 1/2-Rényi divergence | Preserves the probabilistic interpretation of FLN weights; true metric; gently penalizes differences by comparing square roots of probabilities; reveals biologically plausible subdivisions; linear scaling with interareal distance | Interpretation less familiar than correlation-based measures |
| Weighted Jaccard ($J_{\mathcal{W}}$) | Min–max overlap of nonnegative vectors | Incorporates weights; reduces to $J_{\mathcal{S}}$ in binary case | Underestimates overlap for probability mass functions [8]; grouped F1 with somatosensory areas (3,2) |
| Jaccard probability ($J_{\mathcal{P}}$) | Extension of Jaccard to probability vectors; normalizes overlap probabilistically | Preserves probability properties; close match to $H^2$ partitions; reveals premotor–motor subdivision | Cluster area 1 with 3-2 rather than a mixed membership with motor areas; produces higher absolute dissimilarities than $H^2$ |
| Cosine similarity | Angular separation between vectors | Handles signed weights; widely used; related to correlation | Produced biologically inconsistent clusters (e.g., grouped 7m/V6 with V1–V4; F1 with V1–V2); does not respect probabilistic interpretation |
| Pearson correlation | Cosine similarity of $z$-scored vectors | Captures relative deviations from mean connectivity | Similar issues to cosine: grouped dorsal with ventral visual areas; placed F1 with early visual cortex; biologically less plausible |

the professors (nodes) are "performing research in the field of physics and subfield of condensed matter". In another group, other professors are "performing research in the field of physics and subfield of particle physics," etc. In the chemistry department, a similar structure holds, with different research groups. Within each group there are numerus back-and-forth interactions (frequent similar interactions). There are fewer interactions between the research groups, and those are typically at faculty meetings where decisions are made affecting all groups. In this case, from the point of view of the department chair, these are groups "performing research in physics," only – on a higher level of the functional hierarchy. The nature of discussions during faculty meetings will be specific to physics-based research needs. In the chemistry department, the same process happens and at their faculty meetings the discussions (interactions, functional links) will be specific to chemistry-based research needs. The two types of discussions in the two departments will clearly be different, although they will have commonalities. The two departments form two different nodal communities. The next level is at the College of Science, which houses (inclusion) both departments (and others). The dean is a mediator between these departments and when the dean has a meeting of the chairs (each representing their departments), the discussions are at the level of departmental research needs and the potential return on those research activities (and less on research group specifics within the departments). When an issue arises that is beyond a particular research area (such as research space availability, graduate student stipends, etc.) then these functional activities (links) are running between all the departments, because decisions here globally impact the College of Science. At the next level, that of the provost, similar interactions occur, but now between the deans of the different colleges (Science, Arts and Letters, Engineering, etc.) through links that affect all at that level. If there was only links going in one direction between two groups than those groups do not belong to the same level of the hierarchy, one is subordinate to the other.

##### S3.3 Identifying the optimal node-community hierarchy level from the optimal link-community hierarchy

We may use the loop entropy to find an optimal link-community hierarchy level. However, finding the associated node-community hierarchy level is not straightforward. In S16, we explain our method with an example.

##### S3.4 Cover assignment algorithm

At a selected level (other than the first) from the node-community hierarchy, there might be single-node communities. However, it is still desired to assign them to one or several non-single node communities, i.e., with more than one member. In the S17 and its caption we explain how the algorithm works, with an illustration.

##### S3.5 Omega index

It is a measure of the agreement between community partitions of a network when nodes can belong to more than one group or cover. To the best of our knowledge, it was introduced in [4].

Assume there is a network with $N$ nodes partitioned into two cover structures

$$C = \{C_i\}_{i=1}^{k} \quad \text{and} \quad C' = \{C'_{i'}\}_{i'=1}^{k'},$$

such that $C_i$ ($C'_{i'}$) denotes the set of nodes in the $i$-th ($i'$-th) cover, which are not necessarily mutually exclusive. Then, their omega index is defined as

$$\omega(C, C') = \frac{\omega_u(C, C') - \omega_e(C, C')}{1 - \omega_e(C, C')},$$

where

$$\omega_u(C, C') = \binom{N}{2}^{-2} \sum_{j=0}^{\max(k,k')} \left|t_j(C) \cap t_j(C')\right|,$$

and

$$\omega_e(C, C') = \binom{N}{2}^{-2} \sum_{j=0}^{\max(k,k')} \left|t_j(C)\right| \left|t_j(C')\right|.$$

In the above equations, $t_j(\cdot)$ represents the set of node pairs that appear exactly $j$ times in a cover. For example, suppose there are four nodes $a, b, c, d$ with two cover structures

$$C = \{\{a, b, c\}, \{c, d\}\} \quad \text{and} \quad C' = \{\{a, b\}, \{c, d\}, \{a, d\}\},$$

with two and three covers, respectively. Then $j \in \{0, 1, 2, 3\}$. For $j = 0$,

$$t_0(C) = \{(a, d), (b, d)\}, \qquad t_0(C') = \{(a, c), (b, d), (b, c)\}.$$

Thus, $|t_0(C) \cap t_0(C')| = 1$, meaning that only the pair $(b, d)$ appears zero times in both cover structures. Note that if a node pair belongs to $t_0$, it cannot belong to any other $t_j$ for $j > 0$; hence the sets $t_j(C)$ are disjoint.

In this formulation, $\omega_u$ represents the probability of randomly selecting a node pair with equal frequency of occurrence in both cover structures for some $t_j(\cdot)$, whereas $\omega_e$ denotes the expected probability of such agreement under independence. The normalization factor $1 - \omega_e$ ensures that $\omega = 1$ if and only if the node-pair frequency sets $t_j(\cdot)$ are identical across both cover structures for all $j$, denoting perfect similarity or overlap.

#### S3.6 Performance analysis

To measure the performance of our algorithm, we generated weighted and directed Lancichinetti–Fortunato (LF) benchmarks with ground-truth node community memberships, allowing some nodes to participate in more than one cover [6] (see S14). The benchmark generates networks with power-law degree and power-law community-size distributions. Here, we choose the degree and community size distribution exponents to be $t_1 = 2$ and $t_2 = 1$, respectively.

Additionally, the benchmark has a topological mixing parameter $\mu_t$ and a weight mixing parameter $\mu_w$ that control the complexity of the generated networks. Suppose $k_i$ is the total degree of node $i$, and define the total strength of the node as $s_i \equiv k_i^{\beta}$, with $\beta = 2$. The internal degree $k_i^{(\text{in})}$ and internal strength $s_i^{(\text{in})}$ are the number of neighbors of node $i$ and the sum of the weights of the links between node $i$ and all nodes in the same cover as node $i$. They are defined as

$$k_i^{(\text{in})} \equiv (1 - \mu_t) k_i, \qquad s_i^{(\text{in})} \equiv (1 - \mu_w) s_i,$$

with $\mu_t, \mu_w \in [0, 1]$. By definition, the external degree $k_i^{(\text{ext})}$ and external strength $s_i^{(\text{ext})}$ are

$$k_i^{(\text{ext})} = k_i - k_i^{(\text{in})}, \qquad s_i^{(\text{ext})} = s_i - s_i^{(\text{in})}.$$

Low values of $\mu_t$ and/or $\mu_w$ imply that most of the links or link strengths of a node remain within its own cover, indicating a sharp community structure. In contrast, large values of $\mu_t$ or $\mu_w$ correspond to a more diffuse community structure.

Let $w_{ij}$ denote the weight of the link from node $i$ to node $j$. The set of link weights $\{w_{ij}\}$ is generated randomly and then adjusted to minimize

$$\text{Var}(\{w_{ij}\}) = \sum_i \left[ (s_i - \rho_i)^2 + \left( s_i^{(\text{in})} - \rho_i^{(\text{in})} \right)^2 + \left( s_i^{(\text{ext})} - \rho_i^{(\text{ext})} \right)^2 \right],$$

where

$$\rho_i = \sum_j w_{ij}, \qquad \rho_i^{(\text{in})} = \sum_j w_{ij}\, \kappa(i, j), \qquad \rho_i^{(\text{ext})} = \sum_j w_{ij}\, (1 - \kappa(i, j)),$$

and $\kappa(i, j) = 1$ if nodes $i$ and $j$ share at least one community membership, and 0 otherwise.

We varied $\mu_t$ between 0.1 (compact communities; low complexity) and 0.8 (diffuse communities; high complexity) in steps of 7/90 (10 values), while fixing $\mu_w = 0.01$. A low $\mu_w$ was chosen because these networks are sparse; otherwise, the external strength $s^{(\text{ext})}$ would be concentrated in a small number of inter-community links, making them disproportionately strong compared to within-community links, which would be biologically implausible.

Other parameters include the average and maximum degree, $\langle k \rangle \in \{5, 7, 10\}$ and $k_{\max} \in \{30, 50\}$ (S14), and the minimum and maximum community size, set to 5 and 25, respectively. We generated benchmarks with $N = 100$ nodes, of which 5% had overlapping memberships in 2, 3, or 4 communities (denoted Om, shown as columns in S14), while the rest participated in a single community. For each parameter setting, 100 network instances were generated.

After generating the networks, we applied our algorithm to recover the cover structure, including multi-community memberships. This was done by computing the node-community hierarchy (NCh) of the networks and identifying the optimal level in the link-community hierarchy (LCh) using both the maximum average link-community density $\langle D \rangle$ [1] and the loop entropy $S_L$. The average link-community density $\langle D \rangle$ at level $l$ of the LCh, in a network with $M$ links, is defined as

$$\langle D \rangle_l = \sum_{c \in C(l)} D_c \frac{m_c}{M},$$

where $C(l)$ is the set of link communities at level $l$, and $D_c$ is the *effective density* of link community $c \in C(l)$. This is given by

$$D_c = \frac{m_c - (n_c - 1)}{n_c(n_c - 1) - (n_c - 1)} = \frac{m_c - (n_c - 1)}{(n_c - 1)^2},$$

where $m_c$ and $n_c$ are the number of links and nodes in community $c$, respectively. Here, $n_c - 1$ is the number of links in the tree null model of the community, and $n_c(n_c - 1)$ is the maximum possible number of directed links between its nodes.

Once the optimal level is found in the LCh, we identify the corresponding optimal level in the NCh. We then apply the cover assignment algorithm to single-node communities and compute the omega index $\omega$ between the predicted and ground-truth community structures. S14 shows that $S_L$ generally outperforms $\langle D \rangle$, except in very dense communities (bottom row, $\langle k \rangle = 10$ and $k_{\max} = 30$), where local density provides a more suitable score. In all other cases, $S_L$ yields better identification of the optimal node-community partition.

Finally, we note that weighted, directed networks with overlapping community memberships generated by the LF benchmark are highly challenging for most state-of-the-art community detection algorithms. To our knowledge, none of these methods are effective in highly dense networks. In contrast, our algorithm produces a node-community hierarchy that encodes community information across multiple scales.

##### S3.7 Using interareal distance as a dissimilarity measure

Here we briefly consider an alternative to the Hellinger distance (which is ultimately based on connectivity strengths), namely the physical interareal separation distance as a dissimilarity measure. At first sight, this seems reasonable, since areas that are more dissimilar in function are often also further apart in space. Moreover, according to S11**b**, $D_{1/2}$ scales linearly, on average, with the interareal distance. However, there is also a substantial amount of variation around this average, which likely contains valuable information not captured by the mean relationship. Indeed, this is the case, as shown in S18 and described in more detail in its caption.

#### S4 Supplementary Figures

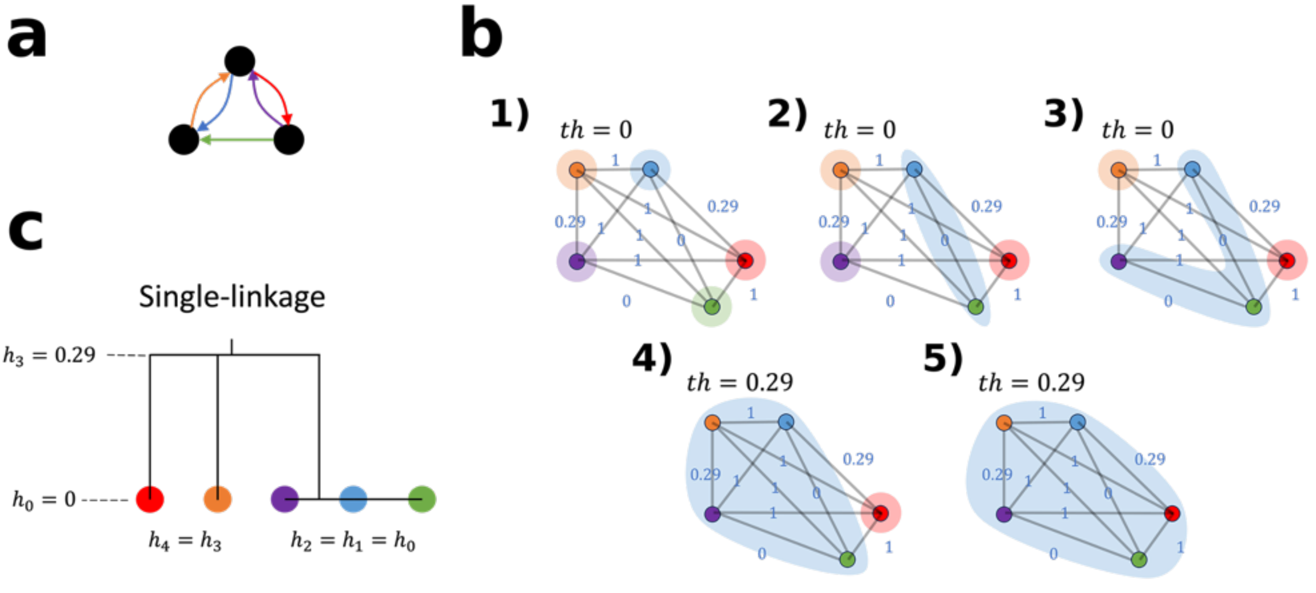

Figure S8: **Link community algorithm example.** **a**, To illustrate how the link community algorithm works, we use a toy binary directed network with three nodes and five links. Each link is colored differently. **b**, The network's nodes are associated with the links in **a**. The $H^2$ dissimilarity between links is shown in blue. **b**1–**b**3 The algorithm starts by setting the lowest link dissimilarity threshold $th = 0$, then merging pairwise those links whose dissimilarity equals the threshold. **b**4, If no more link community pairs can be merged, the threshold is increased to the next lowest dissimilarity value. Note that as soon as the dissimilarity between two members (one from each link community) equals $th$, the link communities are merged. This rule is known as the single-linkage criterion. **b**5, The process of merging link community pairs and increasing the threshold continues until only one link community remains. **c**, The information about which link community pairs merged at each $th$ value is stored in the link-community hierarchy.

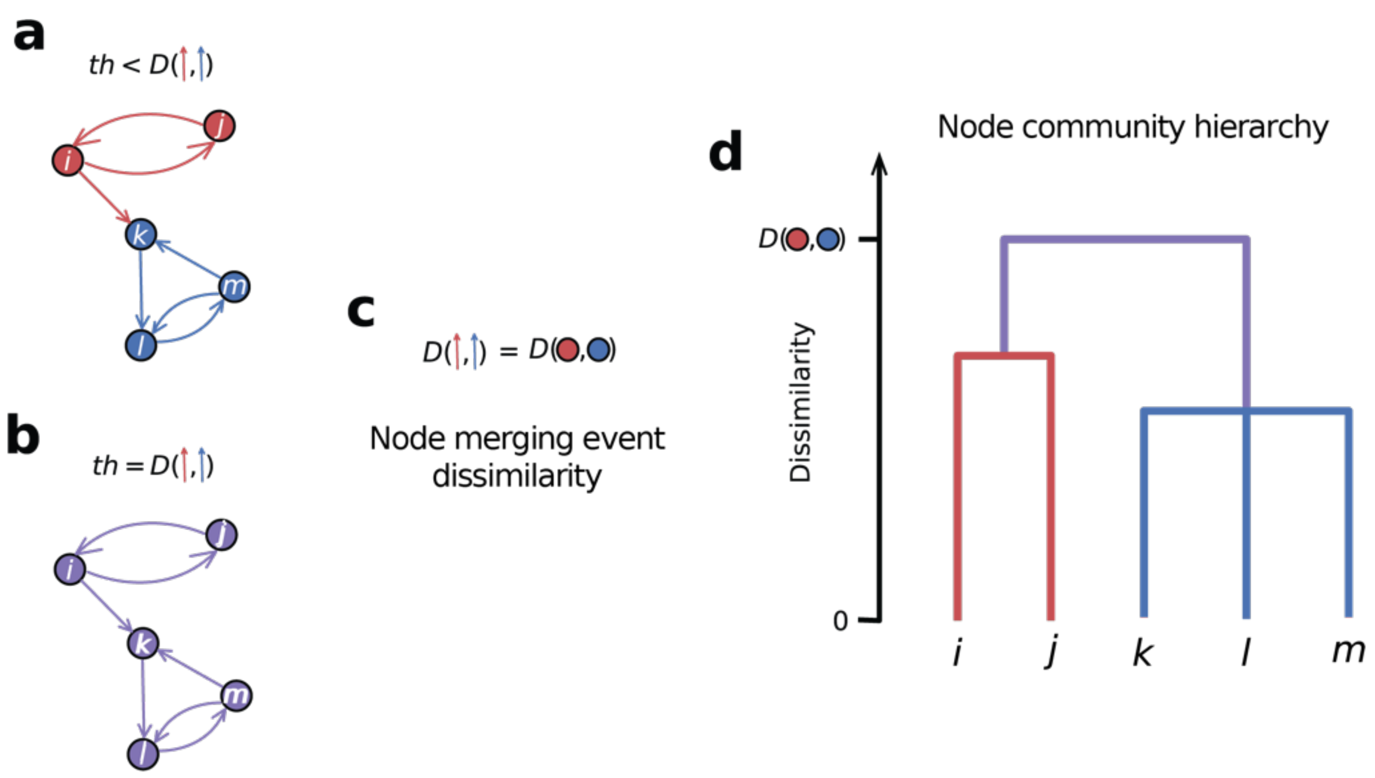

Figure S9: **Workflow for constructing the node-community hierarchy.** **a**, Toy network with 5 nodes and 7 links. At this stage of the link-community hierarchy, the threshold $th$ is smaller "enough" than the distance $D$ between the red and blue link communities (indicated by the arrows), so the two link communities remain disjoint. The corresponding node communities are shown with the same color scheme. **b**, When the threshold reaches $th = D$, the red and blue link communities merge into a purple link community. By the node merging criterion, the associated red and blue node communities also merge into a single purple node community. **c**, Although the link community algorithm does not define distances between node communities, we track node community merging events using the dissimilarity inherited from their corresponding link communities. Here, the distance $D$ between the red and blue link communities is transferred as the dissimilarity between the merged node communities. **d**, Dendrogram of the node-community hierarchy. Coarse-grained node communities emerge as node communities merge. The merging of the red and blue communities into the purple community is represented by a purple branch at height $D$, reflecting the dissimilarity inherited from the link-community hierarchy. This encodes the hierarchical emergence of node communities across link community partitions.

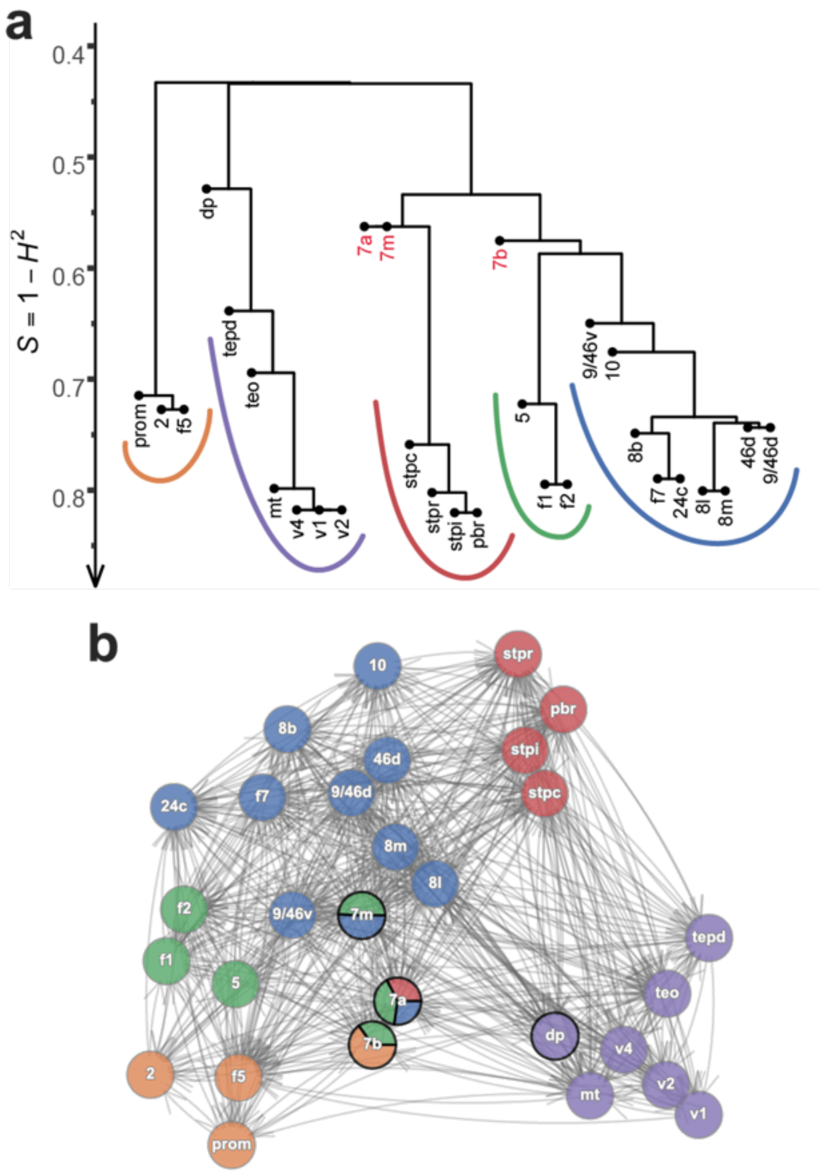

Figure S10: **Community analysis of the edge-complete network of a previous FLN macaque dataset with 29 injected areas** [7]. **a**, All branches of this node-community hierarchy correspond to one branch of Figure **4**d. This demonstrates that the hierarchical community structure of the interareal cortical network is robust, since 11 additional injections do not substantially alter the structure already observed with 29. Red areas denote the NOCs shown in panel **b**. **b**, At the optimal loop-entropy partition, there are five node communities and three NOCs. Only area 7b is also classified as a NOC in the edge-complete subgraph with 40 injected areas. As expected, NOCs are more sensitive to network changes. However, some are conserved and persist even in the final dataset, after all areas have been injected.

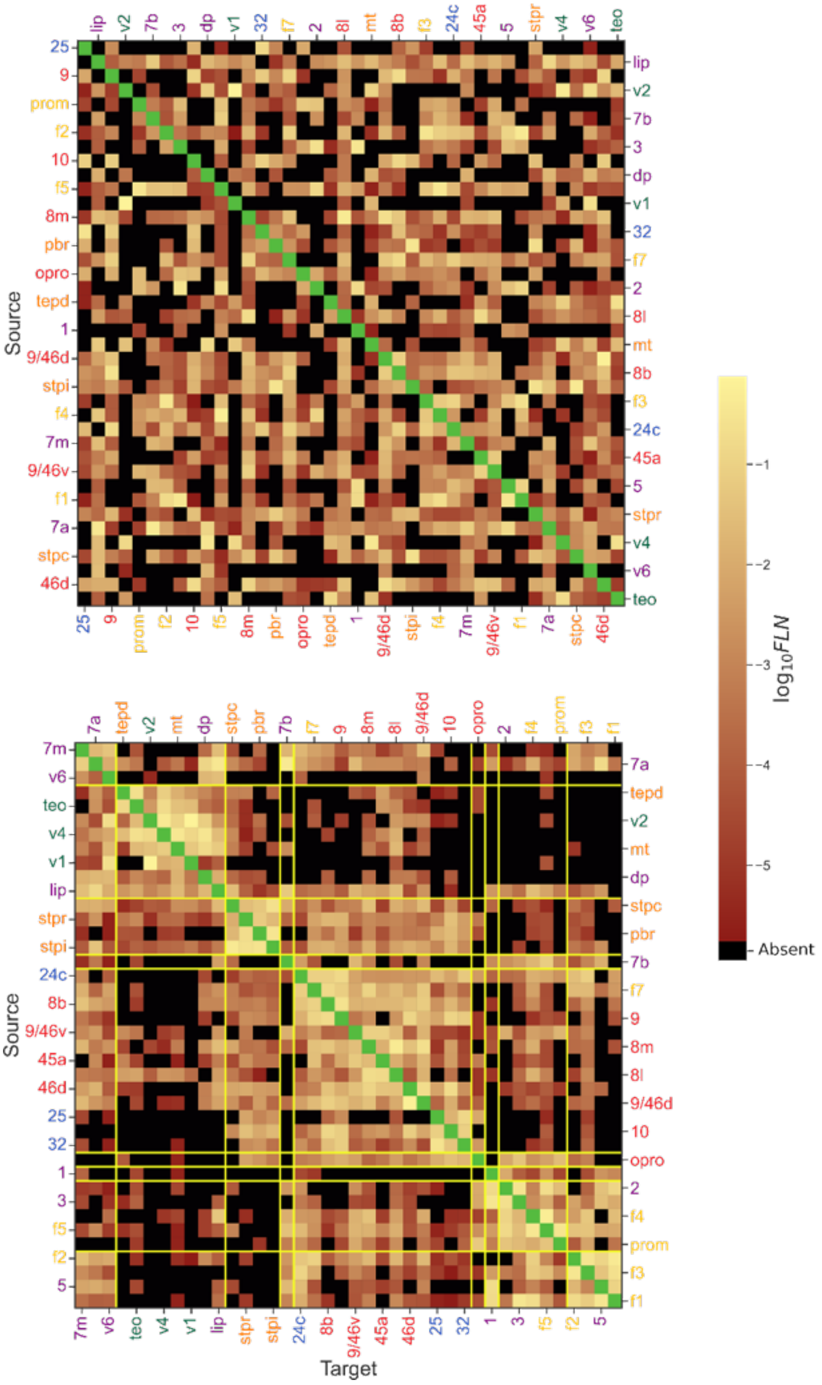

Figure S11: **Connection strength heatmaps of the edge-complete graph.** (**Top**) Rows and columns arranged in random order. (**Bottom**) Rows and columns ordered according to the node-community hierarchy. Yellow vertical and horizontal lines indicate the borders between node communities at the optimal loop entropy level. Absent and intrinsic connections are shown in black and green, respectively. Without information about the network's community structure, the graph is difficult to distinguish from a structureless random network.

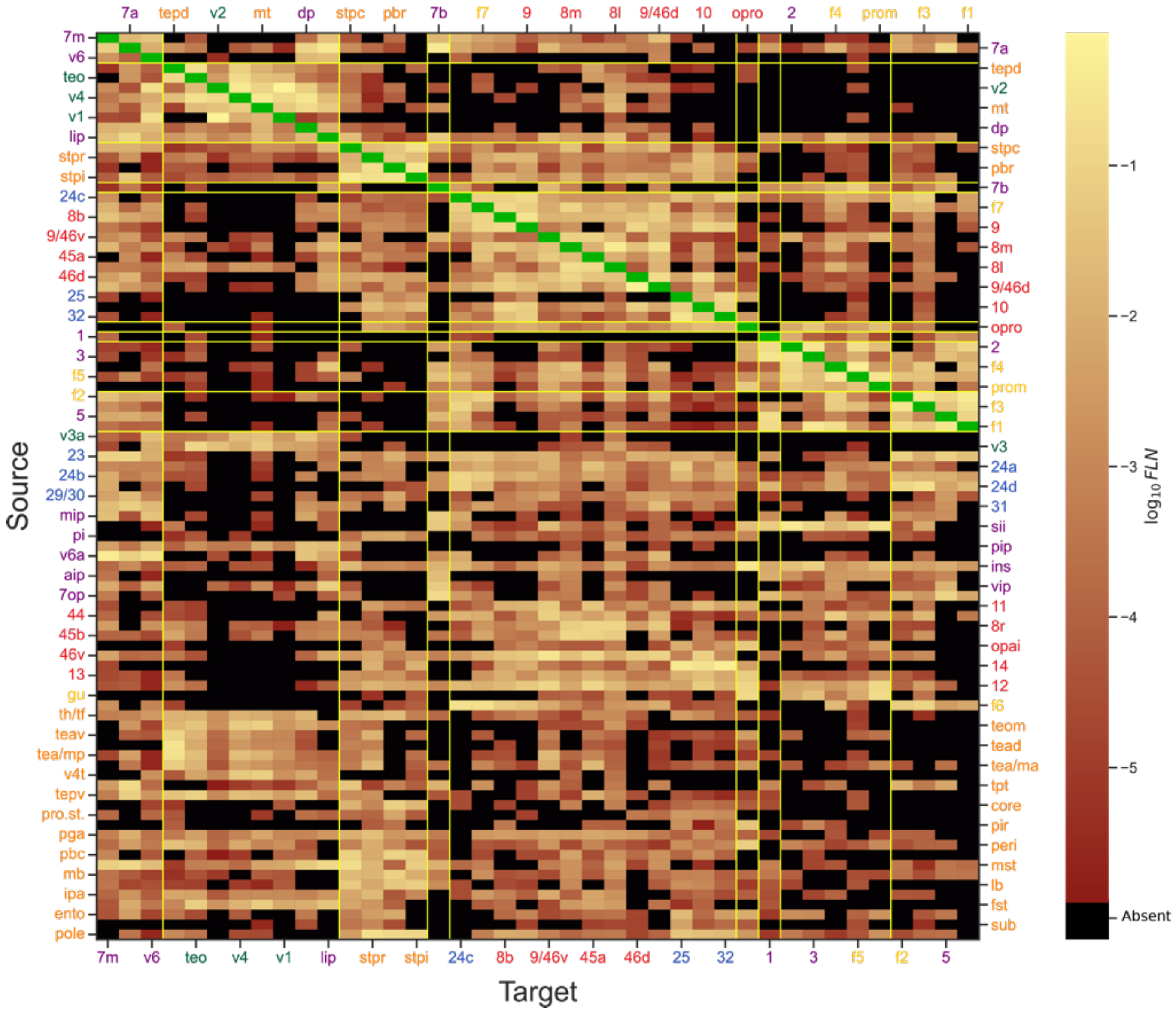

Figure S12: **Sorting of the** $91 \times 40$ **FLN network by communities from the** $40 \times 40$ **subgraph.** Shown is the weighted adjacency matrix, where rows represent out-neighborhoods and columns represent in-neighborhoods of each area. The ordering highlights alignment within and between communities: areas with similar connectivity profiles cluster together, producing visible block structures along the diagonal. The contrast between communities arises from differences in connectivity patterns, seen as variations across rows and columns. Rows below the 40th correspond to non-injected areas, which are sorted by region.

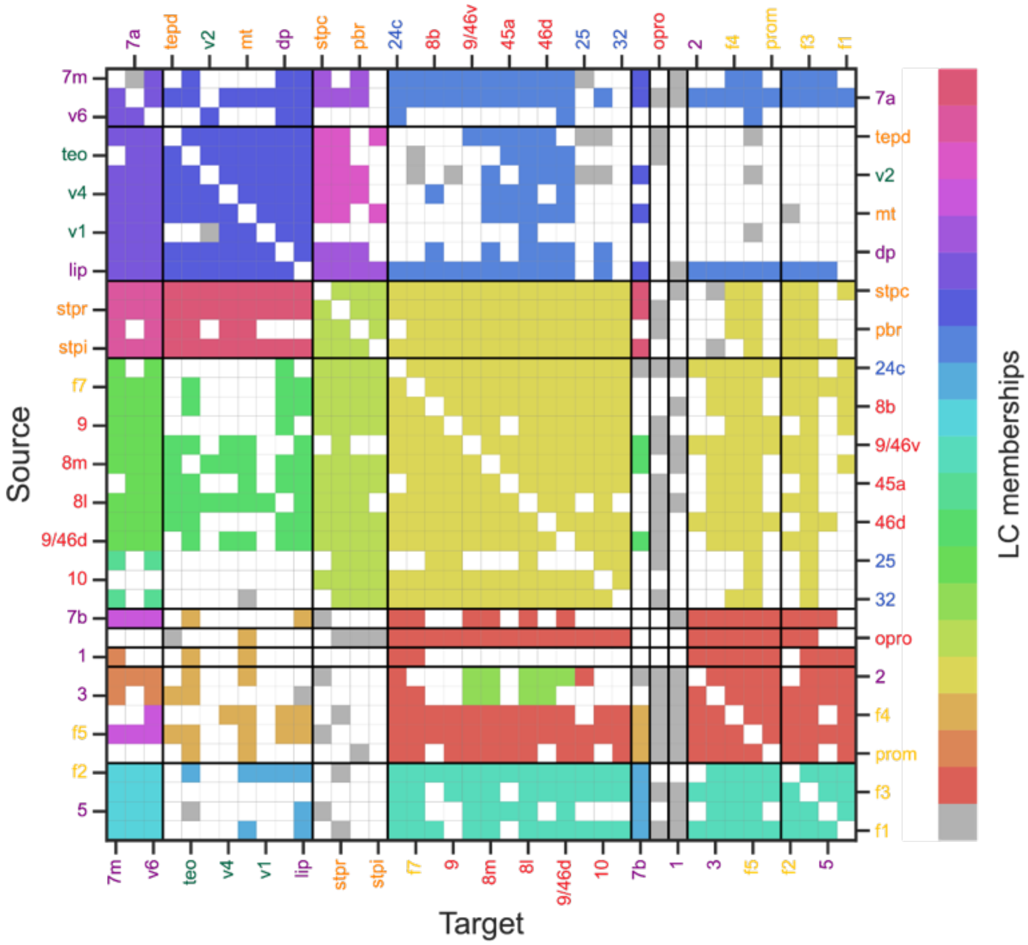

Figure S13: **Link communities at the optimal level.** Links are colored according to their link community membership, with gray links indicating membership in tree link communities. At the maximum loop entropy level, there are 56 link communities, 20 of which correspond to non-tree link communities (non-gray). Rows and columns follow the same ordering as in Figure S11. Using our methodology, link communities tend to connect nodes within the same node community or between specific node communities, a key feature that enhances their interpretability. In addition, links associated with single-node communities are typically tree links, reflecting high dissimilarity with other network links and possibly following distinct organizational principles.

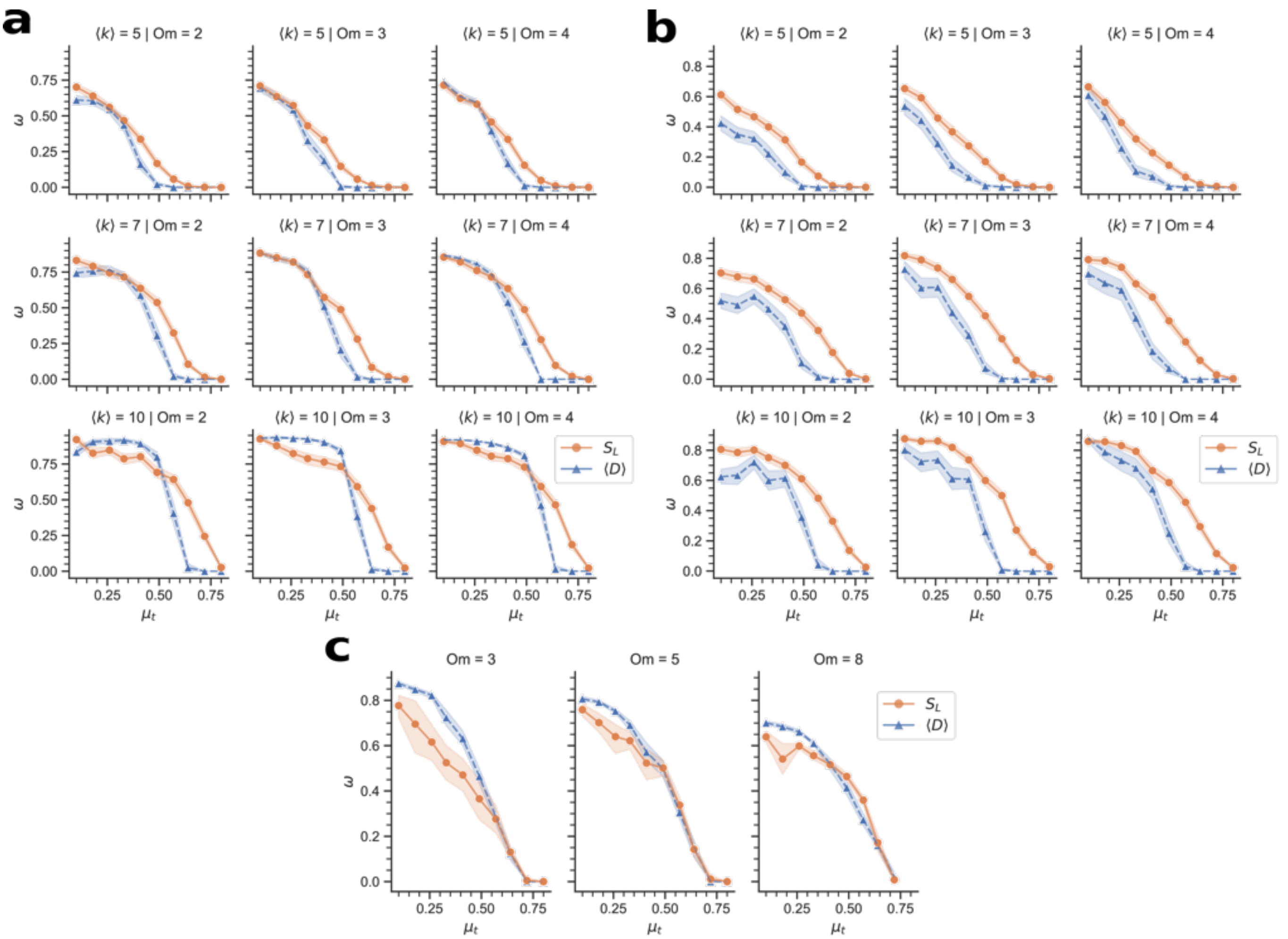

Figure S14: **Algorithm validation.** **a–b**, We used the LF benchmark [6] to generate weighted and directed random networks with ground-truth community structure, allowing some nodes to belong to more than one community. Each network has 100 nodes, with 5% assigned to 2, 3, or 4 overlapping community memberships (Om; columns). The average degree $\langle k \rangle$ is 5, 7, or 10 (rows). We varied the topology mixing parameter $\mu_t$ from 0.1 to 0.8 in steps of 7/90 to assess performance across networks of increasing complexity. Panels **a** and **b** correspond to networks with maximum node degree of 30 and 50, respectively. For each parameter setting, we generated 100 instances, computed the node-community hierarchy, identified the optimal level using the average link community density $\langle D \rangle$ and loop entropy $S_L$, assigned covers to any single-community nodes, and compared the predicted and ground-truth cover structures using the omega index $\omega$. The analysis shows that $S_L$ tends to outperform $\langle D \rangle$ in most cases, except in the bottom row of panel **a**, where the combination of low average degree and low maximum degree yields sparse networks with few external links; in this case, local density is a more suitable community score. Markers and shaded areas indicate averages and one standard deviation, respectively. **c**, Case with $\langle k \rangle = 10$, maximum degree 50, and 1000 nodes, with 10 iterations per $\mu_t$.

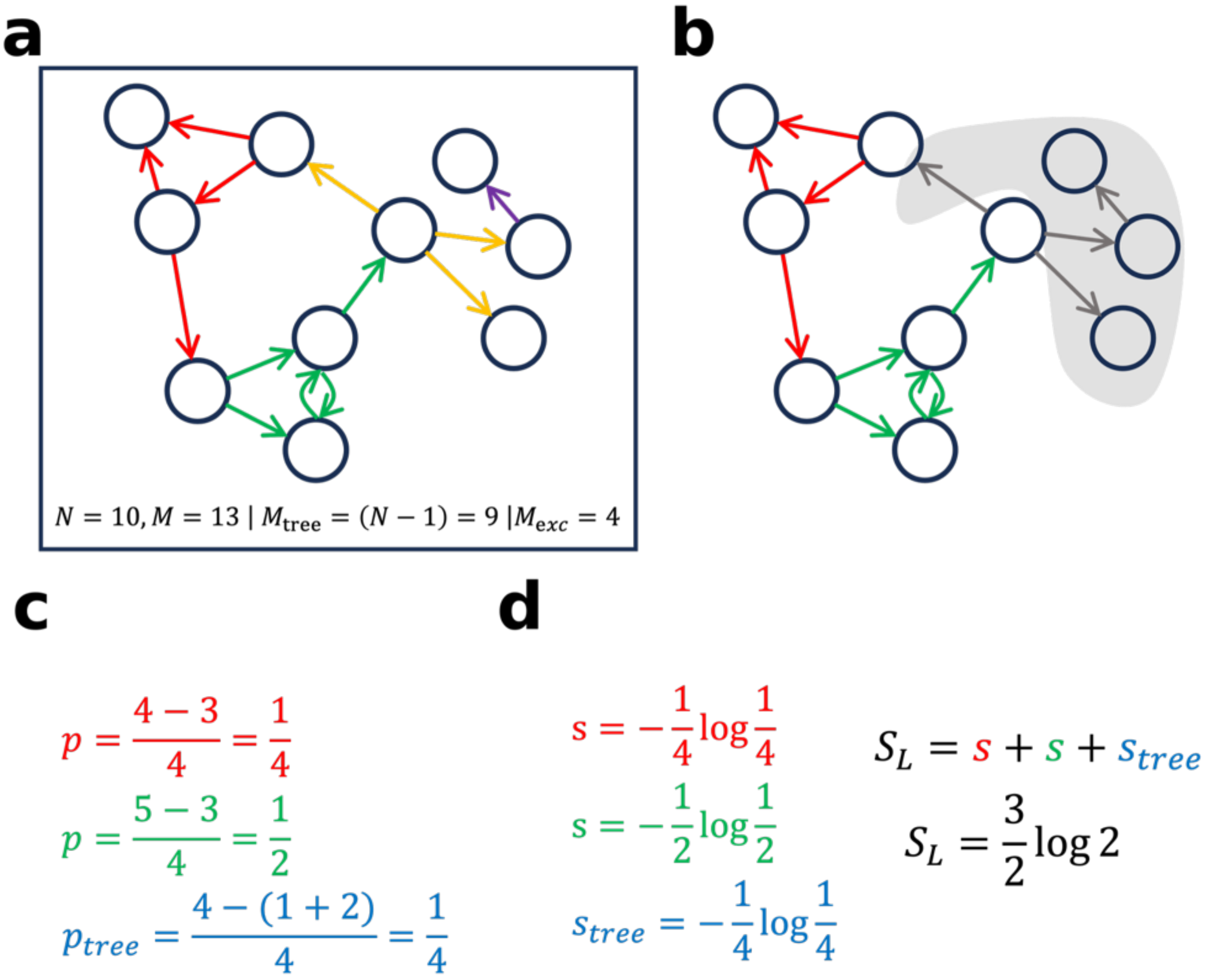

Figure S15: **Loop entropy example.** **a**, Toy network with 10 nodes and 13 directed links. Links are distributed across five link communities. The number of excess links $M_{\text{exc}}$ in this network is four. **b**, The red and green link communities are the only ones that are not trees, since $M_{\text{exc}} > 0$. **c**, Computation of the probability of sampling excess links in the red and green link communities, together with the normalization factor $p_{\text{tree}}$. **d**, The loop entropy $S_L$ is then defined as the Shannon–Gibbs entropy of the excess-link probability distribution.

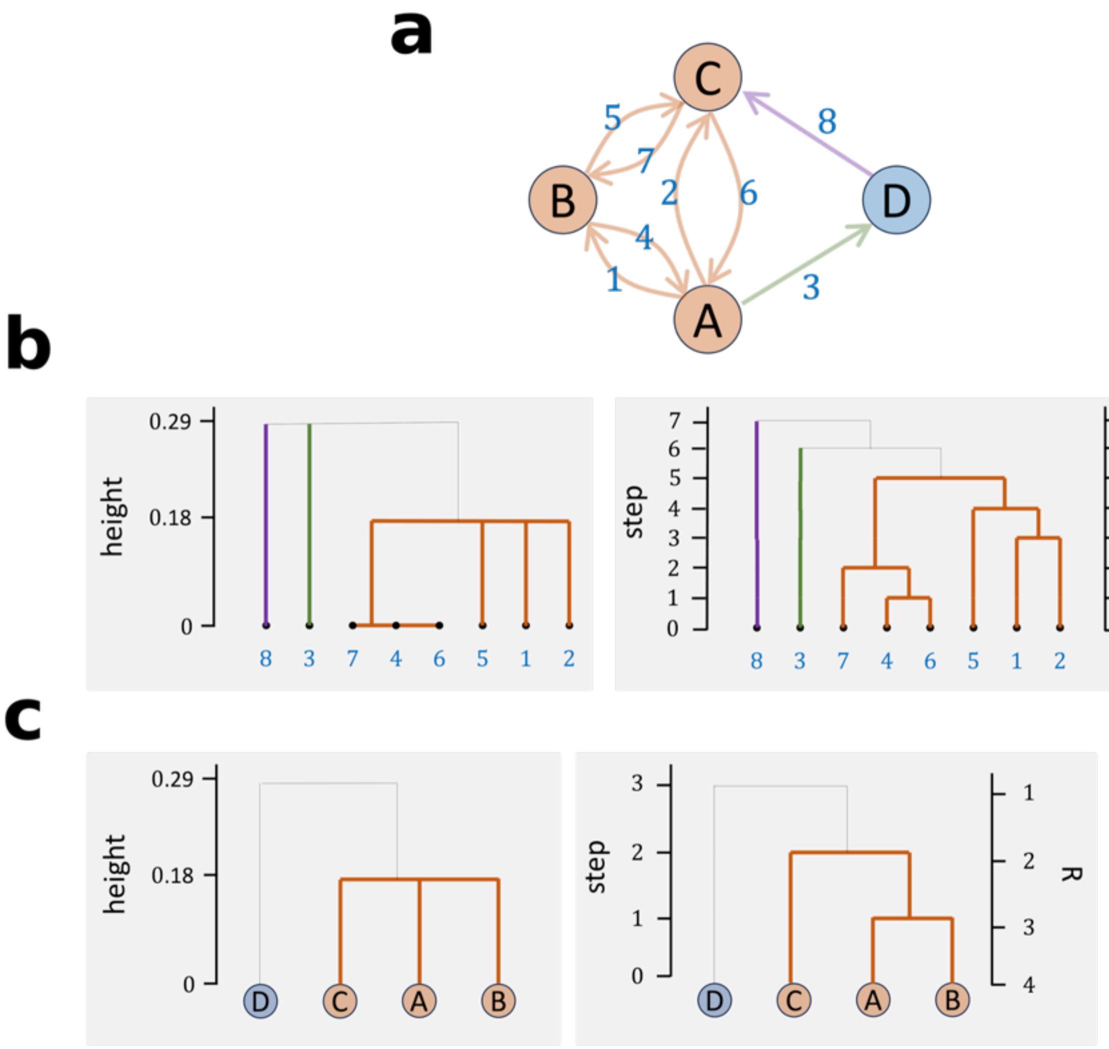

| K | R | height |
|---|---|---|
| 8 | 4 | 0 |
| 7 | 4 | 0 |
| 6 | 4 | 0 |
| 5 | 4 | 0.18 |
| 4 | 4 | 0.18 |
| 3 | 3 | 0.18 |
| 3 | 2 | 0.18 |
| 2 | 2 | 0.29 |
| 1 | 1 | 0.29 |

Figure S16: **Finding a node-community hierarchy level from a link-community hierarchy level.** **a**, Toy network with four nodes and eight directed links. Links are numbered, and their colors represent link communities found at the optimal level. Node communities associated with that level are shown in the same colors. Node “D” forms a single-node community. **b**-left, Link-community hierarchy of the network, with levels indicated by threshold (height). **b**-right, Same hierarchy but with levels separated by link-merging steps (left vertical axis) and number of link communities $K$ (right vertical axis). Each leaf node at the bottom corresponds to a link from the toy network. Branches represent the progression of the link-merging process, with thick colored branches indicating distinct link communities. **c**-left, Node-community hierarchy of the same network, with levels separated by threshold (height). **c**-center, Hierarchy levels separated by node-merging steps. **c**-right, Hierarchy levels by number of node communities $R$. Thick colored branches indicate non-single-node communities. By tabulating $K$, $R$, and height across the link-merging process, one can determine how many node communities $R$ exist at any link-community hierarchy level $K$. If multiple node-merging events occur at the same $K$, the one with the smallest $R$ is selected.

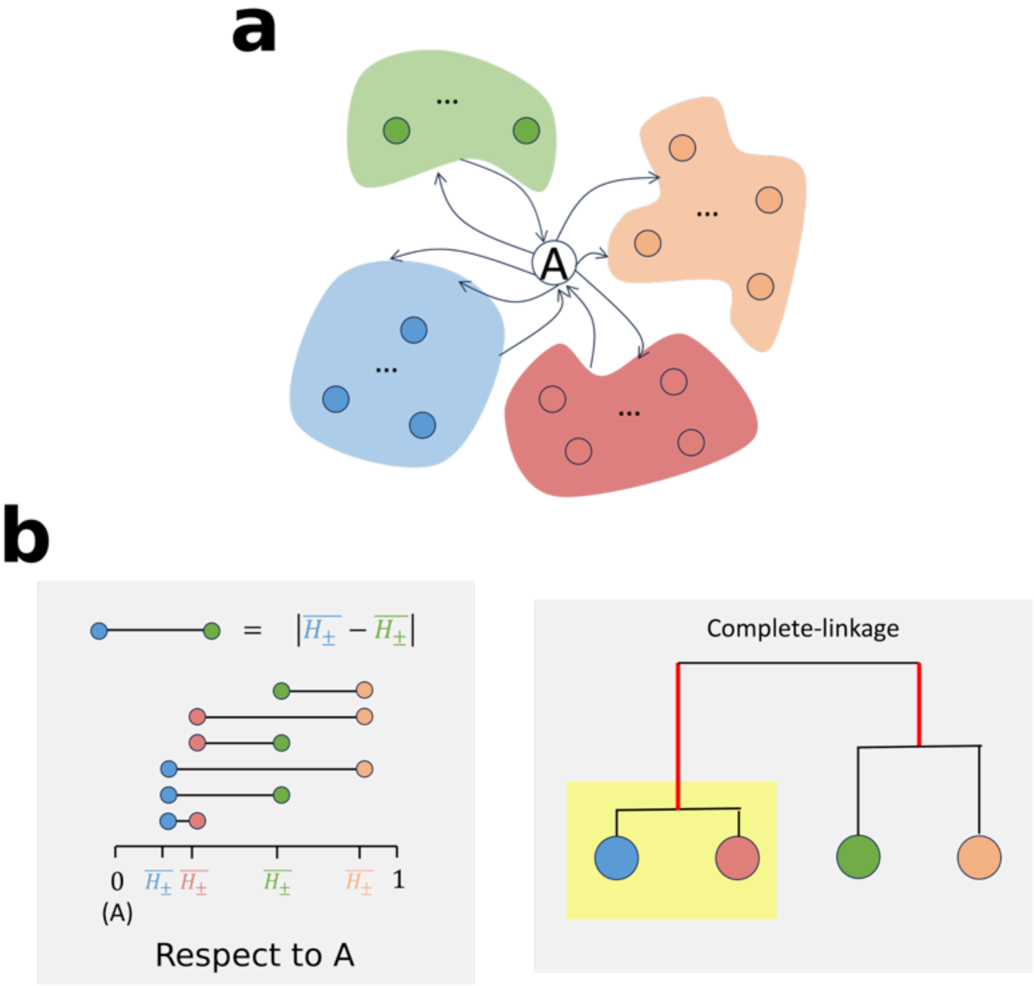

Figure S17: **Cover assignment algorithm.** **a**, Illustration of a single-node community: node “A” does not share a community with any other node in the network. We aim to assign it to one or more of the existing non-single-node communities, shown by colored shapes. **b**-left, For each directed neighborhood, we compute the mean Hellinger distance $\overline{H_{\pm}}$ between the single-node community and the nodes in each non-single-node community. This distance defines the dissimilarity centroid (DC) between the single-node and a non-single-node community. Since $H$ is a metric, the relative differences between $\overline{H_{\pm}}$ values also satisfy the triangle inequality. **b**-right, For each direction, the DCs are clustered using hierarchical agglomerative clustering with the complete-linkage criterion, i.e., the dissimilarity between groups is the maximum dissimilarity between their elements [9]. This creates a DC hierarchy. Complete linkage is chosen because DCs are expected to form highly separable clusters. The hierarchy is then cut at the level just before the largest increase in dissimilarity threshold (thick red branches), and the single-node community inherits the membership(s) of the DCs in the branch containing the most similar centroid.

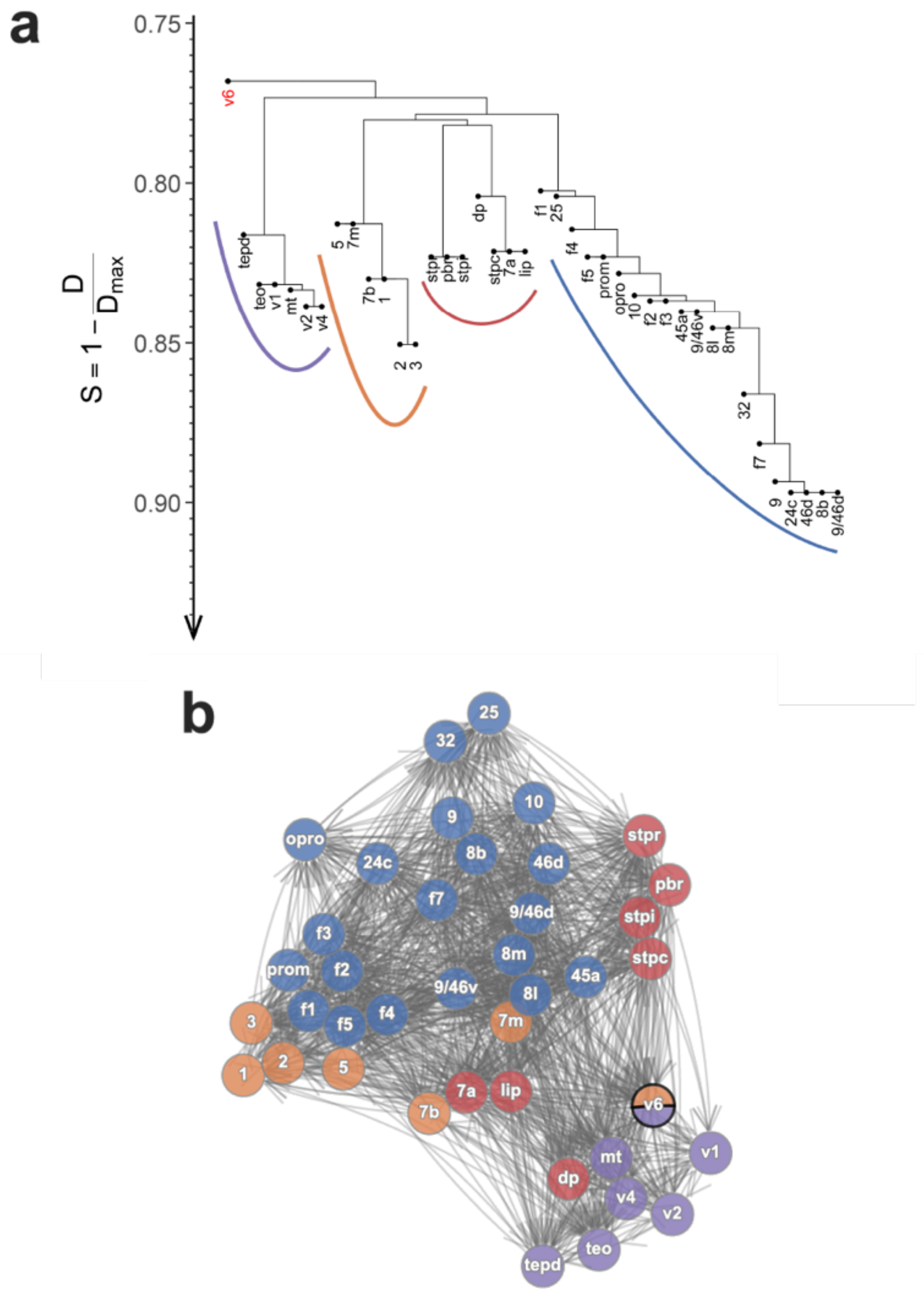

Figure S18: **Community analysis using interareal distance as a dissimilarity measure. a**, Node-community hierarchy obtained by replacing areal similarities with $1 - d/\max(d)$, where $d$ is the interareal distance and $\max(d)$ is the maximum whole-brain interareal distance. Several commonalities and differences arise compared with the hierarchy based on the Hellinger dissimilarity measure (H-NCh). For example, the ventral visual system (above the purple arc) is nearly identical to that in the H-NCh. In contrast, other branches differ substantially. A marked discrepancy occurs in the branch above the blue arc, consisting of prefrontal areas, which shows no sub-branches and thus no evidence of functional specialization. Area V6 (red) is classified as a NOC in this model, in contrast to the H-NCh. **b**, At the optimal loop entropy level, four communities are found, fewer than the six shown in Figure **4**d. The omega index between this partition and the empirical data is $\omega = 0.48$, close to the value for the EDR model ($\omega = 0.44$; see Figure **6**a). This indicates that, apart from the ventral visual system, the interareal distance model does not capture the functional components of the macaque edge-complete subgraph. Overall, the interareal distance-based hierarchy encodes only about 50% of the functional structure recovered with Hellinger distances.
